# Heartbeat-like dynamics drives oxygen activation in methane monooxygenase

**DOI:** 10.1101/2025.01.28.635075

**Authors:** Yunha Hwang, Bumhan Ryu, Dong-Heon Lee, Hyo Jin Hong, Jeong-Geol Na, Chul Gyu Song, Hyun Goo Kang, Edwin Pozharski, Seung Jae Lee

**Author notes:** These authors contributed equally to this work (Yunha Hwang, Bumhan Ryu).

## Abstract

Soluble methane monooxygenase (sMMO) is an enzyme that hydroxylates methane (CH_4_), a potent greenhouse gas, at non-heme di-iron active sites under atmospheric conditions. The regulatory component (MMOB) is essential for the catalytic activity of hydroxylase (MMOH) as it induces conformational changes in the active site and facilitating substrate ingress. Recent advances in cryogenic electron microscopy (cryo-EM) have enabled us to elucidate the high resolution picture of sMMO catalytic mechanism. We describe the 2.85 Å cryo-EM structure of MMOH–MMOB, with one equivalent of MMOB bound to MMOH (H-1B), which is in contrast with previously solved crystal structures. MMOB allosterically regulates the MMOH protomer (αβγ) and induces conformational changes that propagate from the surface to the di-iron coordination site. The *N*-terminal region of the MMOH β-subunit (NT-Hβ) stabilizes helices essential for iron coordination and oxygen activation. The MMOB-bound protomer (HB^A^, αβγB) presents the first structural report of a 2.7 Å Fe···Fe distance, while the non-MMOB-bound protomer (HB^B^, αβγ) and MMOH display a 3.1 Å distance. The coordination of Fe–ligands is maintained by the structural stabilization provided by the β- and γ-subunits of MMOH. This novel cryo-EM structure reveals new coordination environments, offering crucial mechanistic insights into sMMO catalysis.

## MAIN BODY

Carbon dioxide (CO_2_) is a major greenhouse gas, but methane (CH_4_), while generated in lower amounts, has a disproportionate effect due to its ability to persist for longer periods (*1–4*). Methane monooxygenases (MMOs) from methanotrophic bacteria, including sMMO and particulate MMO, can convert methane to methanol under ambient conditions (*5–7*). Understanding the catalytic mechanisms of their action can provide a basis for developing catalysts aimed at reducing atmospheric methane. sMMO belongs to the bacterial multi-component monooxygenase (BMM) superfamily, which includes the key hydroxylase (MMOH) and regulatory (MMOB) components (*1, 8, 9*). Crystal structures of MMOH, MMOH–MMOB, and MMOH–MMOD (inhibitory component) complexes have been reported previously, providing a foundation for mechanistic studies of O_2_ intermediates and C–H activation by sMMOs (*1, 10–15*). It has been discovered that MMOB and MMOD bind to “canyon regions” of MMOH at the dimeric interface of each protomer (αβγ) and induce conformational changes (*1, 8, 11*). These interactions affect the coordination of di-iron active sites and substrate delivery pathways to accelerate or decelerate the catalytic cycle.

MMOH and its complexes present difficult targets for crystal structure determination owing to their large size and dynamic ability to associate or dissociate with other components (*16–18*). Recent advances in single-particle cryo-EM have allowed us, for the first time, to delineate the structural and dynamic details of these heterogeneous protein complexes, thereby overcoming the limitations imposed by the crystalline environment (*19–21*). Being a BMM superfamily member makes sMMO a suitable candidate for understanding its mechanisms using cryo-EM. We investigated MMOH and MMOH–MMOB from *Methylosinus sporium* 5 to explain atomic details from the near-native structure (*11, 22*). Specifically, the 2.85 Å cryo-EM structure of MMOH–MMOB showed that the complex generated with one molecule of MMOB bound to MMOH (H-1B, PDB: 8XIW) was in stark contrast to the crystal structures which showed two MMOB molecules bound to MMOH (H-2B) as shown in **Fig. 1, fig. S1,** and **Table S1**. The 3D variability analysis (3DVA) of cryo-EM data indicated some presence of two molecules of MMOB bound to MMOH (H-2B), but only in the context of stacking multiple MMOH units (*23–25*) (**Fig. 1A** and **fig. S2**). We also determined a 2.64 Å cryo-EM structure of MMOH (PDB: 8YRD) for comparison to crystalline and complex structures (**fig. S3** and **Table S1**). Resolution enhancement in these structures was achieved by employing particle rebalancing to address preferential orientations and confirmational flexibility (*21, 24–26*) (**Fig. 1** and **fig. S1)**. Structural analysis revealed modes of protein dynamics and a heart-shaped MMOH molecule exhibiting “beating heart” type motions. These results imply that binding of MMOB induces conformational changes, which in turn activate the di-iron catalytic center.

**Fig. 1.**
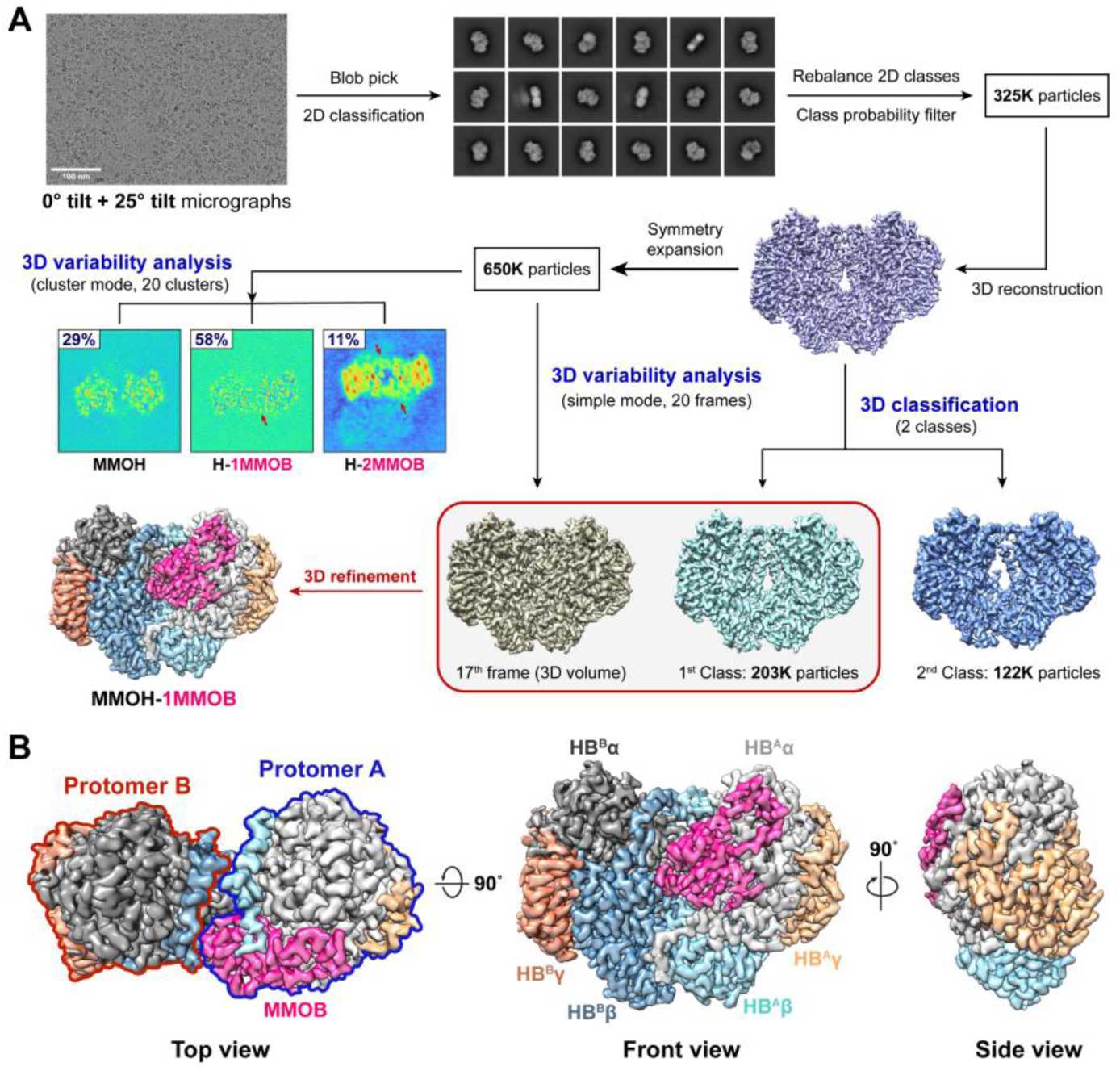
Complex formation of MMOH and MMOB in near-native state. (**A**) Flow chart for cryo-EM image processing for the complexes of MMOH and MMOB (H-1B, PDB: 8XIW). ’K’ denotes a value in thousands. Red arrows depict MMOB in 3DVA with cluster mode. (**B**) Top, front, and side views of the H-1B complex. MMOB (hot pink) binds to the protomer A (α-subunit, gray; β-subunit, aquamarine; γ-subunit, bright orange) of MMOH (HB^A^).

The asymmetric nature of the H-1B assembly provides a window into the mechanism of allosteric activation (**Fig. 1B**). The structural changes in the MMOB-bound protomer (HB^A^, αβγB) and non-MMOB-bound protomer (HB^B^, αβγ) were significantly different from those observed in H-2B crystal structures (*1, 8*) (**fig. S4**). The earlier structures were the most energetically stable forms *in crystal*, which may differ from the vitrified solution state in cryo-EM (*13, 27*). It is possible that while one protomer binds to MMOB to facilitate methane hydroxylation, the other may be interacting with the reductase (MMOR) responsible for electron transfer. Since crystal lattice constraints are more likely to induce a decoy conformation, we propose that the H-1B form of the protein complex is catalytically active, which is further supported by the fact that it produces catalytically competent geometry of the di-iron site (*vide infra*).

Root-mean-square deviation (RMSD) analysis revealed significant conformational differences between MMOH and H-1B. Notably, the *N*- and *C*-terminal regions of each subunit from both structures offer novel insights that were previously unresolved in crystal structures (*1, 10, 28*). The flexible *N*-terminal region of the HB^A^ β-subunit (NT-HB^A^β), which includes Pro4– Thr10, was rigidified upon MMOB binding (**Figs. 2B-C** and **figs. S5A-D**). Structural changes induced by MMOB binding were visualized via 3DVA using particle stacks that included both HB^A^ and HB^B^ protomers (*23, 24*) (**Fig. 1A** and **fig. S1**). This showed significant changes occurring to helices F, G, H, and 4 in HB^A^α; however, NT-HB^A^β was shown to stabilize helices B and C (**Fig 2C** **and fig. S6**). Gln5 and Thr10 of HB^A^β formed hydrogen bonds with Asp71 and Glu222 of MMOB and the HB^A^ α-subunit (HB^A^α), respectively, as shown in **Fig. 2D**. This caused positional shifting of the four-helix bundle, including helices E and F, which affected crucial sidechains involved in substrate delivery and iron coordination, such as Glu243 and Glu209 (**Fig. 2D** **and figs. S7A-B**).

**Fig. 2.**
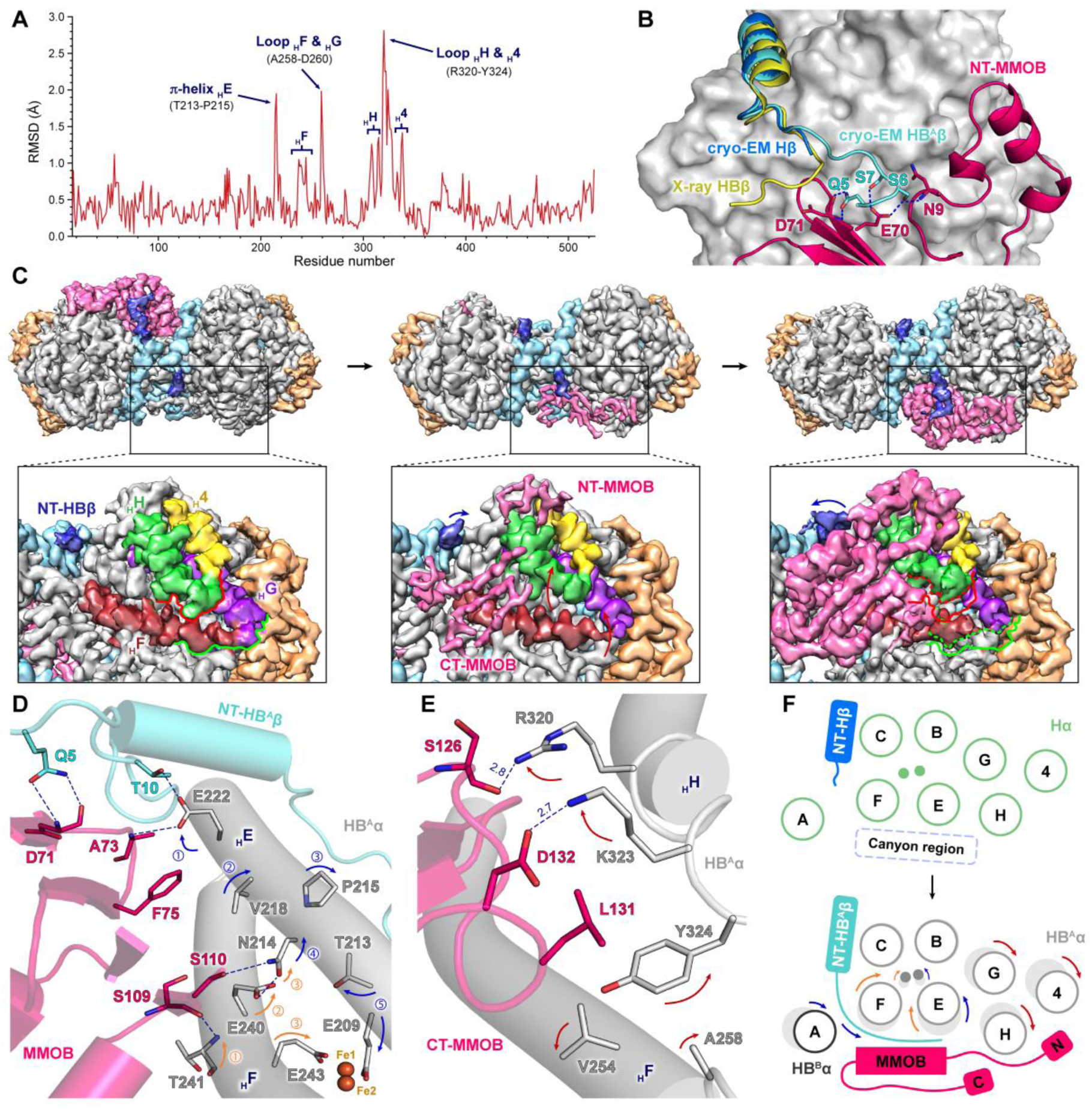
Conformational changes in MMOH induced by MMOB binding. (**A**) Root-mean-square deviation (RMSD) of the main chain (C_α_) per residue for the α-subunit of cryo-EM MMOH and HB^A^. (**B**) Structural alignment of NT-MMOHβ in cryo-EM MMOH (marine), cryo-EM HB^A^ (aquamarine), and X-ray HB (yellow, PDB: 4GAM). Cryo-EM HB^A^α (gray) is shown using a surface representation model. (**C**) Molecular motions of MMOH and MMOB by 3DVA in the simple mode (4^th^, 9^th^, and 18^th^ frame). The black outlined box represents the molecular motion of the MMOH canyon region induced by MMOB binding. The subscript H before an English letter indicates a helix. In the 4^th^ frame, solid lines representing helices F, G, H, and 4 shift upon binding with MMOB, eventually reaching their positions, as represented by dashed lines of the same color in the 18^th^ frame. (**D**) NT-HB^A^β and MMOB affect the di-iron active site by inducing shifts in helices E and F of HB^A^α. (**E**) CT-MMOB induces a shift in helix H of HB^A^α. The color of the arrows indicates conformational changes caused by CT-MMOB (red), NT-HB^A^β (blue), and the core region of MMOB (orange). The unit of distance is Å ngström (Å). (**F**) Schematic model showing the allosteric effect of MMOH by interaction with MMOB.

The *C*-terminal region of MMOB (CT-MMOB) adds two hydrogen bonds via Ser126 and Asp132 and hydrophobic interactions between Leu131 from MMOB and aliphatic and aromatic residues from helices H and F in HB^A^α (**Fig. 2E** and **figs. S7C-D**), which has not been observed in crystal structures (*1*). The allosteric effect of the MMOB docking could be traced to the movement of the long- and short-helices in HB^A^α (**Fig. 2F**). MMOB directly induced movements in helices A, E, F, H, and 4, influencing the positions of helix G (**Fig. 2F** and **figs. S5E-G**), while NT-HB^A^β stabilized helices B and C. These structural changes result in O_2_ activation via first-sphere coordination changes in the di-iron active site, including shortening of the Fe···Fe distance (*vide infra*) (*28–30*).

Binding of the single MMOB molecule to MMOH triggered a cascade of changes from the third sphere to the active site (*1, 8*). Importantly, the π-helical region of helix E transforms to α-helix, driven by sidechain rotations of residues in the four-helix bundle (*31*) (**fig. S4** and **fig. S6**). These movements of helices are key to allosteric activation of the enzyme, facilitating both substrate ingress and electron transfer. The reduced MMOH showed strong binding to the MMOB, and other reports have proposed that MMOB induces a reduced state of MMOH (*32*). Based on the comparison with the crystal structures, binding of the MMOD to the MMOH dimer inhibits the enzyme, which can be explained by additional conformational changes in the four-helix bundle, resulting in reversal of the conformational changes at the active site (*1, 8, 11*). The functions of Hβ were not easily explained, but the H-1B structure proves that *N*-terminal region of the MMOH β-subunit (NT-Hβ) is essential for the catalytic function through the regulation of the four-helix bundle, which contains the active site.

Our study shows that the MMOH γ-subunit (MMOHγ), comprising eight α-helices, is crucial for maintaining structural integrity and enhancing the functional efficiency of MMOH (*10, 28*) (**figs. S8A-C**). Interactions of the MMOHγ with other subunits in MMOH influences the conformational changes necessary for catalytic activity. The precise role of the MMOHγ in regulating enzymatic activity remains poorly understood, highlighting the need for more comprehensive structural and functional studies. The α8-helix located at the *C*-terminal of MMOHγ (CT-Hγ) directly interacts with helices 8 and G to compensate for the translational movement in helices F, G, H, and 4 induced by MMOB binding (**fig. S8D**). Notably, in the cryo-EM H-1B structure, it was observed that the random coil at the *C*-terminal of HBγ (CT-HBγ) aligned alongside the β3-strand of MMOH upon MMOB binding in both protomers, forming a new β-strand (Lys163–Leu167) secondary structure due to the shift of Leu165 in MMOHγ (**figs. S8E-F**). This structural rearrangement stabilized the conformational changes in MMOH induced by the binding of a single MMOB to one side of the canyon region. The core region of MMOHγ interacted with the flexible loops of MMOHβ, enabling the γ-subunit to sustain compression of HB^A^ induced by MMOB binding while allowing relaxation of HB^B^, thereby contributing to the overall structural stability (**figs. S8A-C**). In the MMOH structure itself, Gln55 and Arg116 of MMOHγ form hydrogen bonds with Asp64 and Glu89 of MMOHβ, respectively (**fig. S8G**). Interestingly, upon MMOB binding, these interactions were differentially affected in HB^A^, where the hydrogen bonds were strengthened by compression, whereas in HB^B^, relaxation resulted in the dissociation of these hydrogen bonds (**figs. S8H-I**). This differential interaction highlights the role of the γ-subunit in maintaining structural stability by modulating the conformational states of the two protomers.

MMOB-docking promoted diverse allosteric effects from the surface to the di-iron coordination sites, and HB^A^ demonstrated different coordination from previous reports (**Fig. 3**). In the cryo-EM structure of MMOH, we observed an Fe1···Fe2 distance of 3.1 Å, similar to that of the crystal structure of oxidized MMOH (MMOH_ox_, Fe^3+^–Fe^3+^; **Fig. 3A** and **fig. S9**), which suggests that di-iron is not reduced to di-ferrous by electron irradiation (*1, 8, 28, 30*). The same Fe1···Fe2 distance was observed in the HB^B^ protomer from H-1B, despite some metal ligand coordination changes (**Fig. 3B**). The HB^A^ protomer exhibits dramatic changes that explain enzyme activation (**Fig. 3C**). The Fe1···Fe2 distance from Q-intermediate was reported as 2.70 or 2.84 Å by extended X-ray absorption fine structure and QM/MM calculation (*33, 34*). The Fe1···Fe2 distance decreased to 2.7 Å in HB^A^ due to MMOB binding, because this H-1B complex induced geometric reorganization to generate Q-intermediate, which is in agreement with recent reports (*14*). The Glu144 bidentate coordination to Fe1 and Fe2 was shortened, while the Nδ(His147)-Fe1 distance stretched, increasing from 2.2 Å in MMOH to 2.5 Å in HB^A^ (**Figs. 3B-C** and **fig. S9**). The cryo-EM structure demonstrated enzyme activation from asymmetric binding, providing structural details from a novel di-iron active site. Observing both active and inactive forms of MMOH and comparing these to the non-bound form highlighted conformational changes in the vicinity of the active site triggered by long-range modifications due to MMOB binding. These are summarized in **Table S2**, and the overall results showed compression of the di-iron site and shortening of the Fe1···Fe2 distance, relevant to catalytic activation.

**Fig. 3.**
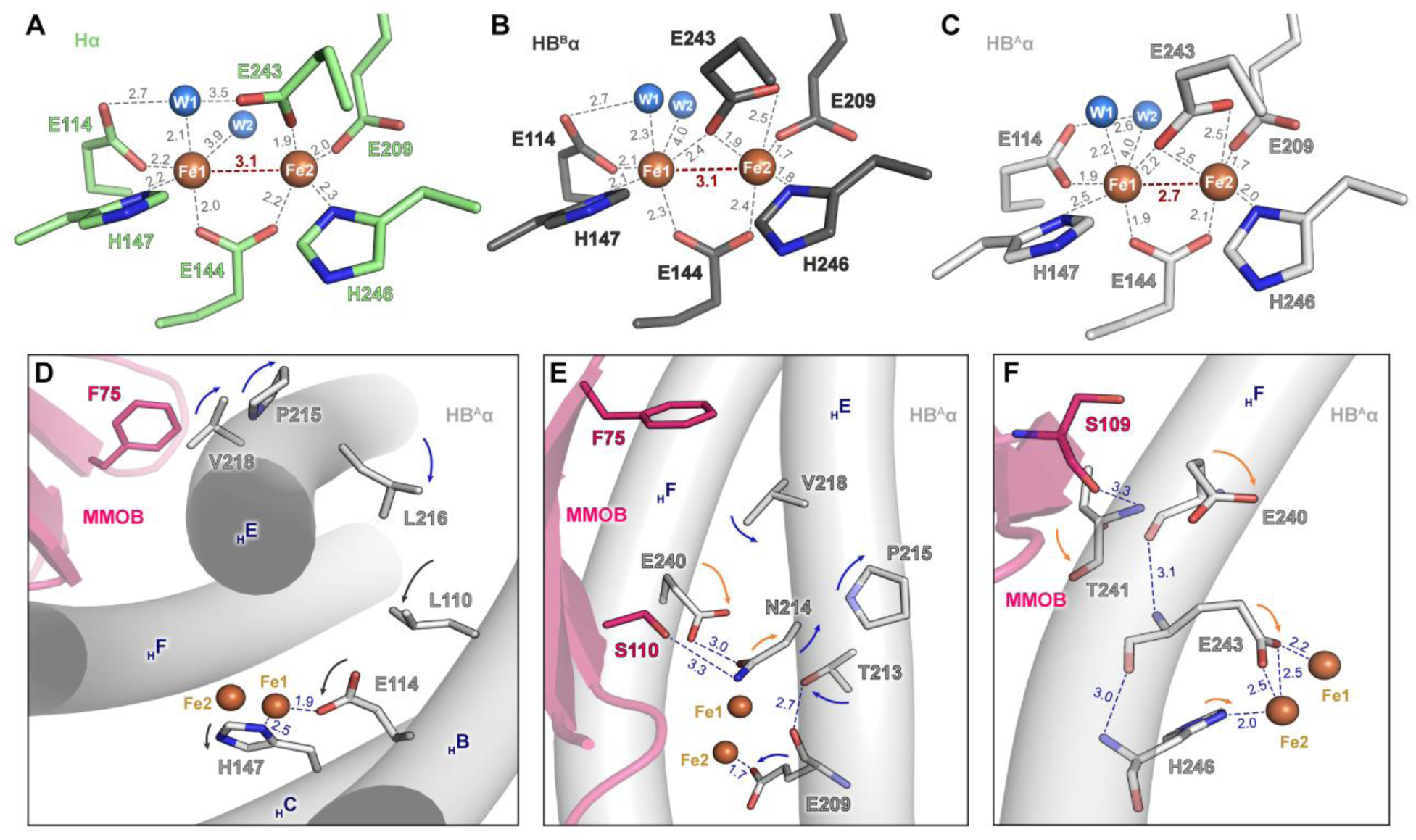
Di-iron active site in MMOH and H-1B from cryo-EM. Geometry of the di-iron center in (**A**) MMOH, (**B**) HB^B^, and (**C**) HB^A^. The two Fe atoms are coordinated by six residues, including four glutamates and two histidines. Water molecules are displayed as marine spheres and are numbered W1 and W2 according to their positions. The coordination distances are depicted in gray, and Fe–Fe distances are represented in red. The unit of distance is Å ngström (Å). (**D**-**F**), F75, S109, and S110 in MMOB induce the shifting of residues in the four-helix bundle, which was related to the coordination of di-iron. The subscript H before an English letter indicates a helix. The color of the arrows indicates conformational changes caused by NT-HB^A^β (blue) and the core region of MMOB (orange). Black arrows indicate conformational changes in the residues of helices B and C.

The electronic and geometric structures of the active site were altered for O_2_ activation via MMOB interaction (*13*). Previous results indicate that MMOB interaction induced a di-ferrous state, while HB^B^ was not photo-reduced (*35, 36*) (**Fig. 3B**). The cryo-EM structure of MMOH coordination shows the enzyme in a typical di-ferric state, despite exposure to potential photo-reduction from the light source (**Fig. 3A**). MMOB docking results in shifting of the carboxylate of Glu243 from Fe2 to Fe1, and the Fe1–Oδ distance is approximately 2.4 Å in HB^B^ (**Fig. 3B**); however, MMOB binding results in compression of the first coordination sphere, shortening the Fe1–Fe2 distance (**Fig. 3C**). Glu243 shows changes of its Fe1 coordination, while the Fe2–Oδ (Glu243) distance increases from 1.9 to 2.5 Å upon MMOB binding (**Figs. 3B-C**). The bidentate coordination of Glu144 approaches di-ferrous upon MMOB binding, while the His147 distance becomes longer (2.1 to 2.5 Å). Glu114 changes its position, with its O-atom losing the hydrogen bond to a water molecule (W1) and forming a new hydrogen bond with another water (W2). Two different protomers from H-1B provide information regarding the native active site, and the coordination environment of HB^B^ could serve as a model compound. MMOB alters the electronic structure by modifying the redox potential in the di-iron site, and its effect on metal coordination provides key details regarding component interactions (*37*).

The modified coordination could be explained by the positional modification of the four-helix bundle triggered by MMOB binding via diverse interactions, including Phe75, Ser110, and Ser109 (**Figs. 3D-F**). Phe75 (MMOB) induces a positional shift of Val218 (HB^A^ α-subunit), and this further shifts Pro215 and Leu216 (helix E) and Leu110 and Glu114 (helix B), as shown in **Fig. 3D** and **figs. S10A-B**. This also induces novel coordination of Glu114 to Fe1. Ser110 and Phe75 in MMOB influences residues such as Val218 and Asn214, which changes the conformation of helix E, and Thr213 generates H-bonds with Glu209 mainchain atoms (**Fig. 3E** and **figs. S10C-D**). These events influence Fe2 coordination. The cryo-EM structure demonstrates that Thr213 acts as a major regulator for significant alterations near the di-iron active site by MMOB, driven by the flexibility of the four-helix bundle. Ser109 (MMOB) forms a hydrogen bond with Thr241 in helix F, and this further changes the Glu240 conformation. These positional modifications finally change the coordination in Glu243 and His246 (**Fig. 3F** and **figs. S10E-F**). Solomon et al. proposed that Fe2 coordination can change in reduced MMOH in the presence of MMOB as supported by circular dichroism (CD), magnetic CD, and Ligand–Field theory of O_2_ reactivity (*38–40*). These studies focused on the disruption of Fe2 by MMOB, and our cryo-EM structure demonstrates that Glu243 and Glu209 are reoriented. Furthermore, MMOB induces electronic and geometric modifications for O_2_ ingress, proving that the modification of Fe1 coordination is necessary to support the reaction intermediates and accelerates the reaction rate (*1, 8*). The H-1B structure shows that MMOB influences both Fe1 and Fe2 coordination.

Possible substrate pathways from the surface to the di-iron active site were investigated, and we find that the cavities 2 and 1 in MMOH_ox_ reported in crystal structures are disconnected (*7, 15*). Previous studies proposed that Phe188 and Leu110 operate as gatekeepers of substrates, but our study suggests that Thr213 also plays a crucial role in substrate access (*1*) (**Fig. 4** and **figs. S11A-F)**. Rotameric alterations in the O-atom from Thr213 causes the hydrophobic cavities to collapse (**Figs. 4A-B** and **figs. S11A-B**). The positional shift in HB^A^ from Leu110 and Thr213 creates hydrophobic cavities in the di-iron active site (**Fig. 4C** and **fig. S11C**). This indicates that MMOB binding optimizes the internal tunnels for O_2_ and substrate access to the di-iron site. The cavities of the HB^B^ protomer were disconnected due to the position of Thr213, which disrupts the hydrophobic pathways to the active site (**Fig. 4B**). The cavities are connected due to the movement of Phe75 from MMOB, which further affects Val218 and Thr213 (**Fig. 3E** and **figs. S10C-D**). The cryo-EM structures of HB^A^ and MMOH indicate that Phe75 induces a hydrophobic shift of Val218, Pro215, and Leu216 from helix E and rotates Leu110 to form hydrophobic substrate-access routes (**figs. S11G-H**). MMOB reorients the positions of the hydrophobic residues in the four-helix bundle, which are crucial for substrate access.

**Fig. 4.**
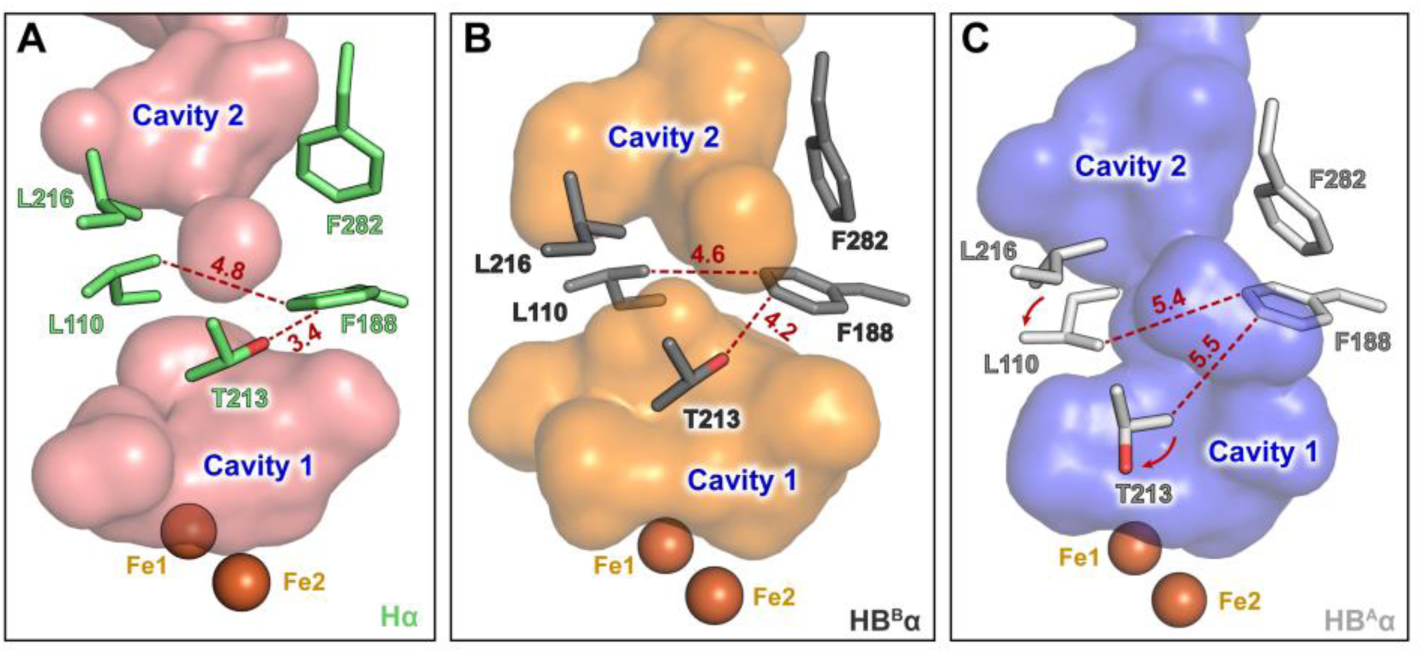
Regulation of the substrate pathway upon MMOB binding to MMOH in the native state. Views of cavities 1 and 2 are shown as translucent van der Waals surfaces in the interior of (**A**) MMOH (residue, lime; surface, deep salmon), (**B**) HB^B^ (residue, dark; surface, orange), and (**C**) HB^A^ (residue, gray; surface, tv_blue). The cavities displayed using PyMOL 2.5.2 are shown as surface models.

To summarize, the cryo-EM structures in this work provide new insights into the activation mechanism of sMMO. The crystal structures solved over a decade ago were constrained by the crystalline environment and likely captured an inhibited conformation of the enzyme. Our study reveals, for the first time, that the active MMOH–MMOB complex is asymmetric, with only a single MMOB molecule bound to the MMOH dimer. This work uncovers a finely tuned allosteric regulation mechanism, wherein conformational changes in the α-helix bundle induce di-iron reorientation and form the substrate access tunnel. These findings significantly advance our understanding of sMMO’s activation and function, setting a new foundation for further studies in enzymatic regulation and methane oxidation.

## Materials and Methods

### Fermentation and purification of MMOH

*Methylosinus sporium* strain 5 MMOH was purified as previously described (*11, 18, 41*). Briefly, MMOH was expressed in *M. sporium* 5 in a copper-limited nitrate minimal salt (NMS) medium at 37 °C in a 5 L incubator and chromatographically purified using DEAE Sepharose Fast Flow (Cytiva), Superdex 200 (Cytiva), and Q-Sepharose Fast Flow (Cytiva) columns attached to the Ä KTA Pure 25 L fast protein liquid chromatography system (Cytiva).

### Expression and purification of MMOB

*M. sporium* 5 MMOB was purified as previously described (*18, 41, 42*). Briefly, MMOB was expressed in *Escherichia coli* BL21(DE3) cells (Novagen) transformed with *mmoB*-pET30a(+) plasmid in Luria Broth media containing kanamycin (50 μg/mL) at 37 °C and chromatographically purified using Q Sepharose Fast Flow (Cytiva) and Superdex 75 (Cytiva) columns attached to the Ä KTA Pure 25 L fast protein liquid chromatography system (Cytiva).

### Cryo-EM grid preparation and data collection

Cryo-EM grids were prepared at the Institute for Basic Science (IBS) and the Institute for Bioscience and Biotechnology Research (IBBR). Purified MMOH and MMOB were stored in buffer C (25 mM MOPS, pH 6.5, 50 mM NaCl, and 1 mM DTT). The MMOH and MMOB (HB) complex was generated by mixing the two proteins in a 1:2.2 molar ratio. MMOH or HB was adjusted to a concentration of 2 mg/mL and applied onto glow-discharged (MMOH, negative; HB, positive) holey carbon grids (Quantifoil, Cu 200 mesh, R 1.2/1.3). After blotting with filter paper, the grid was flash cooled in liquid ethane using Vitrobot Mark IV (Thermo Fisher Scientific; TFS, USA) at 4 °C and 100% relative humidity.

Cryo-EM images were collected at IBS using 300 kV Krios G4 (TFS, USA) equipped with a BioQuantum K3 detector (Gatan Inc, USA) and EPU software (TFS, USA). For MMOH, a total of 8,475 movies were collected. For HB, two separate datasets were obtained from the same sample using two separate grids at different stage angles to overcome the preferred orientation of particles: 3,763 movies at 0° and 11,244 movies at 25°. Further details are provided in **Table S1**.

### Cryo-EM data processing of *M. sporium* 5 MMOH

Data processing was performed using cryoSPARC (v.4.2.1) (*23*). The collected movies were divided into two datasets and pre-processed using patch motion correction and patch contrast transfer function (CTF) estimation. Blob picking from the first dataset produced 4,779,030 particles, which were extracted in a box size of 320 pixels and binned to 256 pixels (resulting in a pixel size of 1.06 Å) (*43*). After two rounds of 2D classification, ab-initio reconstruction generated two 3D classes, one representing the putative protomer (αβγ) and another representing the putative homodimer (α_2_β_2_γ_2_). The particles involved in the putative homodimer were further refined using 3D classification, resulting in the selection of 236,211 particles suitable for the final reconstruction. The resulting particles were re-extracted into 320-pixel boxes and subjected to per-particle refinement jobs (Global and Local CTF refinement). An identical processing workflow was employed for the analysis of the second dataset, and an additional 171,748 particles were acquired. The selected particles from the two datasets were combined and reconstructed with C_2_ symmetry, producing a 3D electron density map at 2.64 Å, where the resolution was estimated via fourier shell correlation at the 0.143 criterion. The dynamics of MMOH were evaluated via 3D flexible training using final particles, and flexible reconstruction was performed (*25*).

### Cryo-EM data processing of the *M. sporium* 5 MMOH–1MMOB complex

Data processing was performed using cryoSPARC (v.3.3.1) (*23*). All movies were pre-processed using Patch motion correction and Patch CTF estimation. Blob pick resulted in 7,714,850 particles, which were extracted in a box size of 360 pixels and binned to 180 pixels (resulting in a pixel size of 1.698 Å) (*43*). After four rounds of 2D classification, oversampled views were removed, which resulted in the selection of 1,535,094 particles (*44*). The resulting particles were re-extracted into 320-pixel boxes. After one round of 2D classification, high-quality particles were re-extracted into 384-pixel boxes and subjected to per-particle refinement jobs (Global and Local CTF refinement). The resulting particles were subjected to 3D variability analysis (3DVA) and 3D classification, respectively (*24*). The 3DVA was performed using 650,028 particles, which resulted from the symmetry expansion of 325,014 particles around the C_2_ symmetry, with a low-pass filter resolution of 3 Å. The 3D variability display was conducted to generate volumes and particles from each of the 20 clusters that fitted the reaction coordinates. The results of this analysis were used to generate linear movies of volumes consisting of 20 frames in simple mode; the 17^th^ frame represented the most stable complex. The particles sorted into the best class were refined using 3D classification, resulting in the selection of 202,837 particles. Homogeneous refinement was performed using imposed C_1_ symmetry with particles selected from 3D classification and volume (17^th^ frame) chosen from 3DVA. One round of 2D classification was conducted with a final selection of 200,043 particles. These were used for 3D refinement with C_1_ symmetry, producing a 3D electron density map at 2.85 Å (estimated from the gold standard fourier shell correlation curve at the 0.143 cutoff).

### Model building and refinement

For the model building of MMOH and the MMOH–1MMOB complex, MMOH_ox_ (PDB: 1MTY) and X-ray MMOH–2MMOB (PDB: 4GAM) structures were fitted into EMD-39540 and EMD-38391 using UCSF Chimera (*1, 28, 45*). The model was enhanced with multiple cycles of real-space refinement in Phenix using geometric and secondary structure restraints, followed by manual corrections in Coot (*46–48*). The overall model validations were performed using the comprehensive validation tool in Phenix (*49*). The statistics of the cryo-EM data collection, 3D reconstruction, and model refinement are summarized in **Table S1**. Structural figures were generated with PyMOL (v.2.5.2) and UCSF Chimera (*45, 50*).

## Acknowledgments

The computational works for this research were performed on the High-Performance Computing Resources at the IBS Research Solution Center.

## Author contributions

Y.H., H.J.H., J.-G.N., and S.J.L. fermented *Methylosinus sporium* 5 and expressed protein; Y.H., C.G.S., and H.G.K. purify the protein; Y.H., B.R., E.P., and S.J.L. conducted electron microscopy; Y.H., B.R., D.-H.K., C.G.S., E.P., and S.J.L. performed electron microscopy image processing and model building; Y.H., D.-H.L., and S.J.L. analyzed coordination studies; E.P. and S.J.L. designed the experiments, compiled the manuscript, and edited and reviewed the experiments and manuscript with input from all the authors.

## Data and materials availability

The data supporting this study are available from the corresponding authors upon request. The cryo-EM structures and the atomic models of the MMOH and MMOH–1MMOB complexes have been deposited in the Electron Microscopy Data Bank (EMDB) and Protein Data Bank (PDB), respectively. The accession codes are as follows: EMD-39540 and PDB ID 8YRD for the cryo-EM structure of *M. sporium* 5 MMOH; and EMD-38391 and PDB ID 8XIW for the cryo-EM structure of *M. sporium* 5 MMOH– 1MMOB complex.

## References and Notes

1. S. J. Lee, M. S. McCormick, S. J. Lippard, U.-S. Cho, Control of substrate access to the active site in methane monooxygenase. Nature 494, 380–384 (2013).

2. S. A. Montzka, E. J. Dlugokencky, J. H. Butler, Non-CO_2_ greenhouse gases and climate change. Nature 476, 43–50 (2011).

3. L. Shen, D. J. Jacob, R. Gautam, M. Omara, T. R. Scarpelli, A. Lorente, D. Zavala-Araiza, X. Lu, Z. Chen, J. Lin, National quantifications of methane emissions from fuel exploitation using high resolution inversions of satellite observations. Nat. Commun. 14, 4948 (2023).

4. J. F. Dean, Old methane and modern climate change. Science 367, 846–848 (2020).

5. F. J. Tucci, A. C. Rosenzweig, Direct Methane Oxidation by Copper-and Iron-Dependent Methane Monooxygenases. Chem. Rev., 124, 1288–1320 (2024).

6. N. F. Dummer, D. J. Willock, Q. He, M. J. Howard, R. J. Lewis, G. Qi, S. H. Taylor, J. Xu, D. Bethell, C. J. Kiely, G. J. Hutchings, Methane oxidation to methanol. Chem. Rev. 123, 6359–6411 (2022).

7. C. E. Tinberg, S. J. Lippard, Dioxygen activation in soluble methane monooxygenase. Acc. Chem. Res. 44, 280–288 (2011).

8. V. Srinivas, R. Banerjee, H. Lebrette, J. C. Jones, O. Aurelius, I.-S. Kim, C. C. Pham, S. Gul, K. D. Sutherlin, A. Bhowmick, J. John, E. Bozkurt, T. Fransson, P. Aller, A. Butryn, I. Bogacz, P. Simon, S. Keable, A. Britz, K. Tono, K. S. Kim, S.-Y. Park, S. J. Lee, J. Park, R. Alonso-Mori, F. D. Fuller, A. Batyuk, A. S. Brewster, U. Bergmann, N. K. Sauter, A. M. Orville, V. K. Yachandra, J. Yano, J. D. Lipscomb, J. Kern, M. Högbom, High-resolution XFEL structure of the soluble methane monooxygenase hydroxylase complex with its regulatory component at ambient temperature in two oxidation states. J. Am. Chem. Soc. 142, 14249–14266 (2020).

9. W. Wang, A. D. Liang, S. J. Lippard, Coupling oxygen consumption with hydrocarbon oxidation in bacterial multicomponent monooxygenases. Acc. Chem. Res. 48, 2632–2639 (2015).

10. A. C. Rosenzweig, C. A. Frederick, S. J. Lippard, P. Nordlund, auml, Crystal structure of a bacterial non-haem iron hydroxylase that catalyses the biological oxidation of methane. Nature 366, 537–543 (1993).

11. H. Kim, S. An, Y. R. Park, H. Jang, H. Yoo, S. H. Park, S. J. Lee, U.-S Cho, MMOD-induced structural changes of hydroxylase in soluble methane monooxygenase. Sci. Adv. 5, eaax0059 (2019).

12. R. Banerjee, Y. Proshlyakov, J. D. Lipscomb, D. A. Proshlyakov, Structure of the key species in the enzymatic oxidation of methane to methanol. Nature 518, 431–434 (2015).

13. C. E. Schulz, R. G. Castillo, D. A. Pantazis, S. DeBeer, F. Neese, Structure–spectroscopy correlations for intermediate Q of soluble methane monooxygenase: insights from QM/MM calculations. J. Am. Chem. Soc. 143, 6560–6577 (2021).

14. R. G. Castillo, R. Banerjee, C. J. Allpress, G. T. Rohde, E. Bill, L. Que Jr., J. D. Lipscomb, S. DeBeer, High-energy-resolution fluorescence-detected X-ray absorption of the Q intermediate of soluble methane monooxygenase. J. Am. Chem. Soc. 139, 18024–18033 (2017).

15. M. Merkx, D. A. Kopp, M. H. Sazinsky, J. L. Blazyk, J. Müller, S. J. Lippard, Dioxygen activation and methane hydroxylation by soluble methane monooxygenase: a tale of two irons and three proteins. Angew. Chem. Int. Ed. 40, 2782–2807 (2001).

16. S. Friedle, E. Reisner, S. J. Lippard, Current challenges of modeling diiron enzyme active sites for dioxygen activation by biomimetic synthetic complexes. Chem. Soc. Rev. 39, 2768–2779 (2010).

17. X. Benjin, L. Ling, Developments, applications, and prospects of cryo-electron microscopy. Prot. Sci. 29, 872–882 (2020).

18. C. Lee, S. C. Ha, Z. Rao, Y. Hwang, D. S. Kim, S. Y. Kim, H. Yoo, C. Yoon, J.-G. Na, J. H. Park, S. J. Lee, Elucidation of the electron transfer environment in the MMOR FAD-binding domain from *Methylosinus sporium* 5. Dalton Trans. 50, 16493–16498 (2021).

19. R. Danev, H. Iijima, M. Matsuzaki, S. Motoki, Fast and accurate defocus modulation for improved tunability of cryo-EM experiments. IUCrJ 7, 566–574 (2020).

20. T. Nakane, A. Kotecha, A. Sente, G. McMullan, S. Masiulis, P. M. G. E. Brown, I. T. Grigoras, L. Malinauskaite, T. Malinauskas, J. Miehling, T. Uchański, L. Yu, D. Karia, E. V. Pechnikova, E. d. Jong, J. Keizer, M. Bischoff, J. McCormack, P. Tiemeijer, S. W. Hardwick, D. Y. Chirgadze, G. Murshudov, A. R. Aricescu, S. H. W. Scheres, Single-particle cryo-EM at atomic resolution. Nature 587, 152–156 (2020).

21. S. Aiyer, P. R. Baldwin, S. M. Tan, Z. Shan, J. Oh, A. Mehrani, M. E. Bowman, G. Louie, D. O. Passos, S. Đorđević-Marquardt, M. Mietzsch, J. A. Hull, S. Hoshika, B. A. Barad, D. A. Grotjahn, R. McKenna, M. Agbandje-McKenna, S. A. Benner, J. A. P. Noel, D. Wang, Y. Zi Tan, D. Lyumkis, Overcoming resolution attenuation during tilted cryo-EM data collection. Nat. Commun. 15, 389 (2024).

22. Y. Hwang, J.-G. Na, S. J. Lee, Transcriptional regulation of soluble methane monooxygenase via enhancer-binding protein derived from *Methylosinus sporium* 5. Applied and Environmental Microbiology 89, e02104–02122 (2023).

23. A. Punjani, J. L. Rubinstein, D. J. Fleet, M. A. Brubaker, cryoSPARC: algorithms for rapid unsupervised cryo-EM structure determination. Nat. Methods 14, 290–296 (2017).

24. A. Punjani, D. J. Fleet, 3D variability analysis: Resolving continuous flexibility and discrete heterogeneity from single particle cryo-EM. J. Struct. Biol. 213, 107702 (2021).

25. A. Punjani, D. J. Fleet, 3DFlex: determining structure and motion of flexible proteins from cryo-EM. Nat. Methods 20, 860–870 (2023).

26. J. Zhu, W. Huang, J. Zhao, L. Huynh, D. J. Taylor, M. E. Harris, Structural and mechanistic basis for recognition of alternative tRNA precursor substrates by bacterial ribonuclease P. Nat. Commun. 13, 5120 (2022).

27. S. J. Amann, D. Keihsler, T. Bodrug, N. G. Brown, D. Haselbach, Frozen in time: analyzing molecular dynamics with time-resolved cryo-EM. Structure 31, 4–19 (2023).

28. A. C. Rosenzweig, H. Brandstetter, D. A. Whittington, P. Nordlund, S. J. Lippard, C. A. Frederick, Crystal structures of the methane monooxygenase hydroxylase from *Methylococcus capsulatus* (Bath): implications for substrate gating and component interactions. *Proteins: Struct., Funct.*, Bioinf. 29, 141–152 (1997).

29. J. García-Nafría, C. G. Tate, Structure determination of GPCRs: cryo-EM compared with X-ray crystallography. Biochem. Soc. Trans. 49, 2345–2355 (2021).

30. D. A. Whittington, S. J. Lippard, Crystal structures of the soluble methane monooxygenase hydroxylase from *Methylococcus capsulatus* (Bath) demonstrating geometrical variability at the dinuclear iron active site. J. Am. Chem. Soc. 123, 827–838 (2001).

31. M. H. Sazinsky, S. J. Lippard, Correlating structure with function in bacterial multicomponent monooxygenases and related diiron proteins. Acc. Chem. Res. 39, 558–566 (2006).

32. W. Wang, S. J. Lippard, Diiron oxidation state control of substrate access to the active site of soluble methane monooxygenase mediated by the regulatory component. J. Am. Chem. Soc. 136, 2244–2247 (2014).

33. D. Rinaldo, D. M. Philipp, S. J. Lippard, R. A. Friesner, Intermediates in dioxygen activation by methane monooxygenase: A QM/MM study. J. Am. Chem. Soc. 129, 3135–3147 (2007).

34. L. M. Dassama, A. Silakov, C. M. Krest, J. C. Calixto, C. Krebs, J. M. Bollinger Jr, M. T. Green, A 2.8 Å Fe–Fe Separation in the Fe_2_^III/IV^ Intermediate, X, from *Escherichia coli* Ribonucleotide Reductase. J. Am. Chem. Soc. 135, 16758–16761 (2013).

35. E. Y. D. Chua, J. H. Mendez, M. Rapp, S. L. Ilca, Y. Z. Tan, K. Maruthi, H. Kuang, C. M. Zimanyi, A. Cheng, E. T. Eng, A. J. Noble, C. S. Potter, B. Carragher, Better, faster, cheaper: recent advances in cryo–electron microscopy. Annu. Rev. Biochem. 91, 1–32 (2022).

36. L. A. Baker, J. L. Rubinstein, Radiation damage in electron cryomicroscopy. Methods Enzymol. 481, 371–388 (2010).

37. B. J. Wallar, J. D. Lipscomb, Methane monooxygenase component B mutants alter the kinetics of steps throughout the catalytic cycle. Biochemistry 40, 2220–2233 (2001).

38. N. a. Mitić, J. K. Schwartz, B. J. Brazeau, J. D. Lipscomb, E. I. Solomon, CD and MCD studies of the effects of component B variant binding on the biferrous active site of methane monooxygenase. Biochemistry 47, 8386–8397 (2008).

39. S. C. Pulver, W. A. Froland, J. D. Lipscomb, E. I. Solomon, Ligand field circular dichroism and magnetic circular dichroism studies of component B and substrate binding to the hydroxylase component of methane monooxygenase. J. Am. Chem. Soc. 119, 387–395 (1997).

40. H. S. Jeong, S. Hong, H. S. Yoo, J. Kim, Y. Kim, C. Yoon, S. J. Lee, S. H. Kim, EPR-derived structures of flavin radical and iron-sulfur clusters from *Methylosinus sporium* 5 reductase. Inorg. Chem. Front. 8, 1279–1289 (2021).

41. Y. R. Park, H. S. Yoo, M. Y. Song, D.-H. Lee, S. J. Lee, Biocatalytic oxidations of substrates through soluble methane monooxygenase from *Methylosinus sporium* 5. Catalysts 8, 582 (2018).

42. C. Lee, Y. Hwang, H. G. Kang, S. J. Lee, Electron transfer to hydroxylase through component interactions in soluble methane monooxygenase. J. Microbiol. Biotechnol. 32, 287 (2022).

43. K. M. Yip, N. Fischer, E. Paknia, A. Chari, H. Stark, Atomic-resolution protein structure determination by cryo-EM. Nature 587, 157–161 (2020).

44. D. M. Snead, M. Matyszewski, A. M. Dickey, Y. X. Lin, A. E. Leschziner, S. L. Reck-Peterson, Structural basis for Parkinson’s disease-linked LRRK2’s binding to microtubules. Nat. Struct. Mol. Biol. 29, 1196–1207 (2022).

45. E. F. Pettersen, T. D. Goddard, C. C. Huang, G. S. Couch, D. M. Greenblatt, E. C. Meng, T. E. Ferrin, UCSF Chimera—a visualization system for exploratory research and analysis. J. Comput. Chem. 25, 1605–1612 (2004).

46. P. V. Afonine, B. K. Poon, R. J. Read, O. V. Sobolev, T. C. Terwilliger, A, Urzhumtsevf, P, D. Adams, Real-space refinement in PHENIX for cryo-EM and crystallography. Acta crystallogr. D Biol. Crystallogr. 74, 531–544 (2018).

47. A. Casañal, B. Lohkamp, P. Emsley, Current developments in Coot for macromolecular model building of Electron Cryo-microscopy and Crystallographic Data. Prot. Sci. 29, 1055–1064 (2020).

48. P. Emsley, K. Cowtan, Coot: model-building tools for molecular graphics. Acta crystallogr. D Biol. Crystallogr. 60, 2126–2132 (2004).

49. P. V. Afonine, B. P. Klaholz, N. W. Moriarty, B. K. Poon, O. V. Sobolev, T. C. Terwilliger, P. D. Adamsa, A. Urzhumtsev, New tools for the analysis and validation of cryo-EM maps and atomic models. Acta crystallogr. D Biol. Crystallogr. 74, 814–840 (2018).

50. W. L. DeLano, Pymol: An open-source molecular graphics tool. CCP4 Newsl. Protein Crystallogr. 40, 82–92 (2002).

