## Supplementary Information for "Heartbeat-like dynamics drives oxygen activation in methane monooxygenase"

**for**

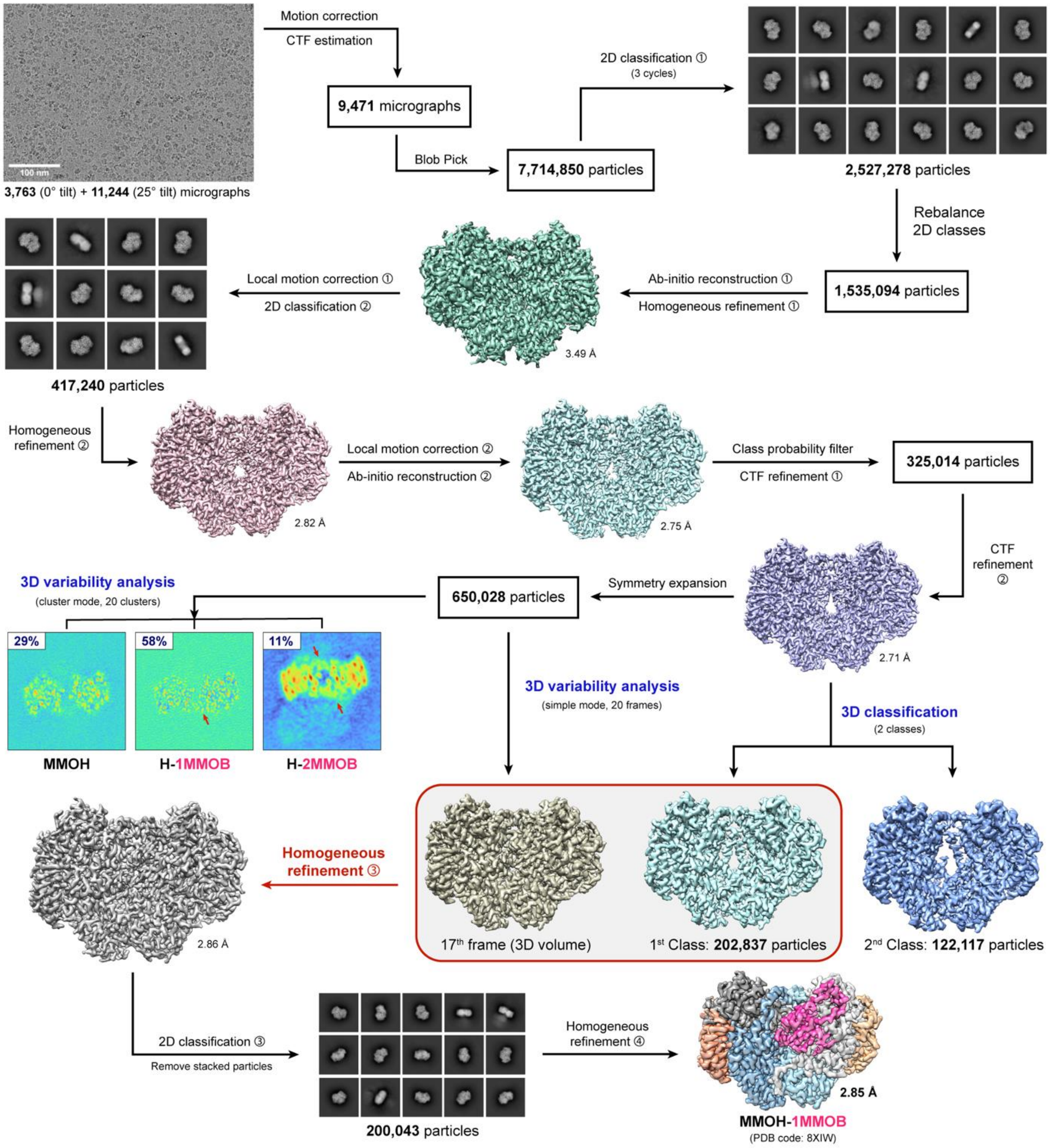


**Fig. S1. Flow chart for cryo-electron microscopy image processing for the complex of MMOH and MMOB (H-1B, PDB: 8XIW).** Red arrows depict MMOB in 3D variability analysis with cluster mode. The number below the 3D map volume indicates the resolution (unit: Å). In the native state, the H-B complex indicated that only one MMOB bound to MMOH.

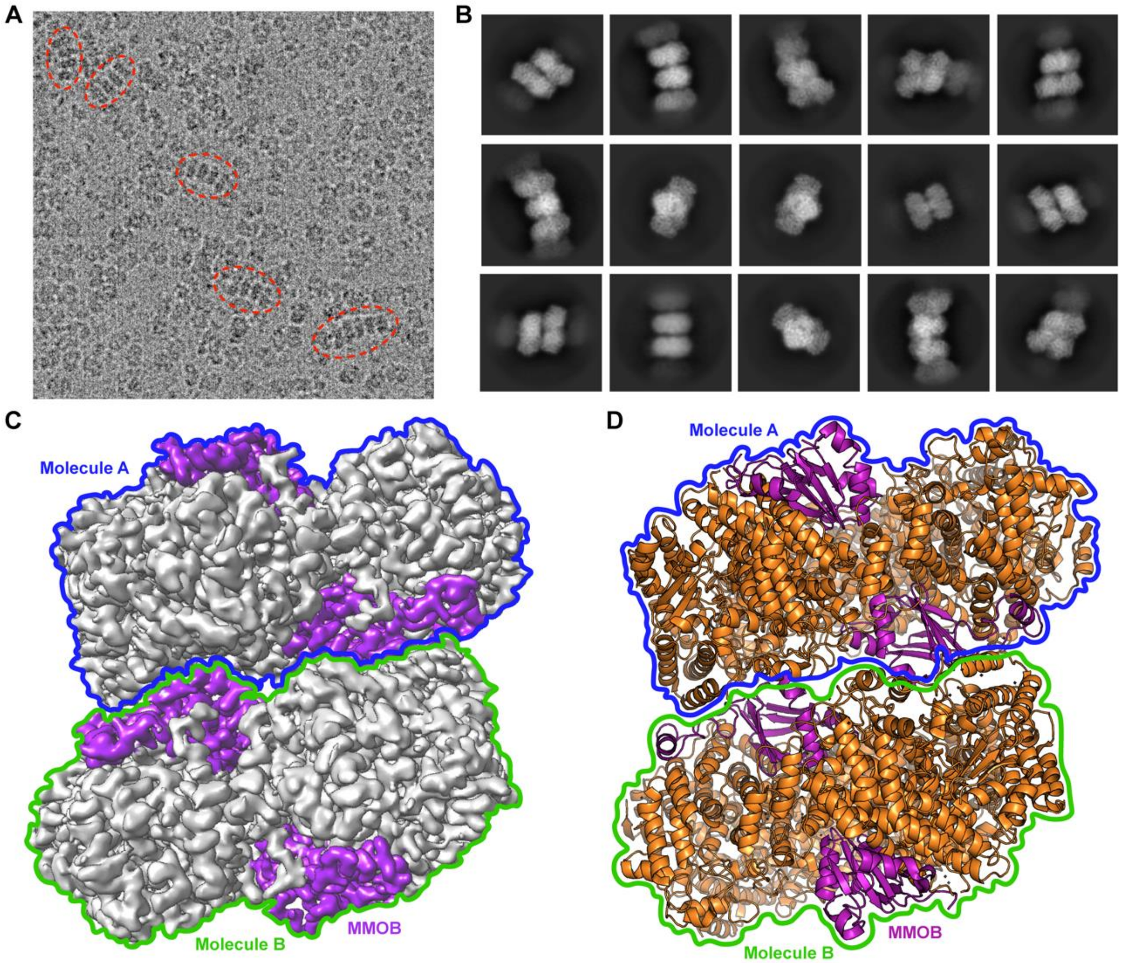


**Fig. S2. H-2B complex from stacked particles of *M. sporium* 5.** (**A**) Stacked particles of the H-2B complex in a micrograph. The H-2B complexes stacked in a row are indicated by red dashed circles. (**B**) The 2D classes of stacked particles of the H-2B complex. (**C** and **D**) Comparison of (**C**) 3D volume of the stacked form about the H-2B complex with cryo-electron microscopy (cryo-EM) and (**D**) X-ray structure (PDB: 4GAM). MMOB is depicted in purple, while MMOH is depicted in gray (cryo-EM) and orange (X-ray crystallography). The stacked MMOH–MMOB complex consisted of two molecules (molecule A, blue; molecule B, green), with each MMOH bound to two MMOB.

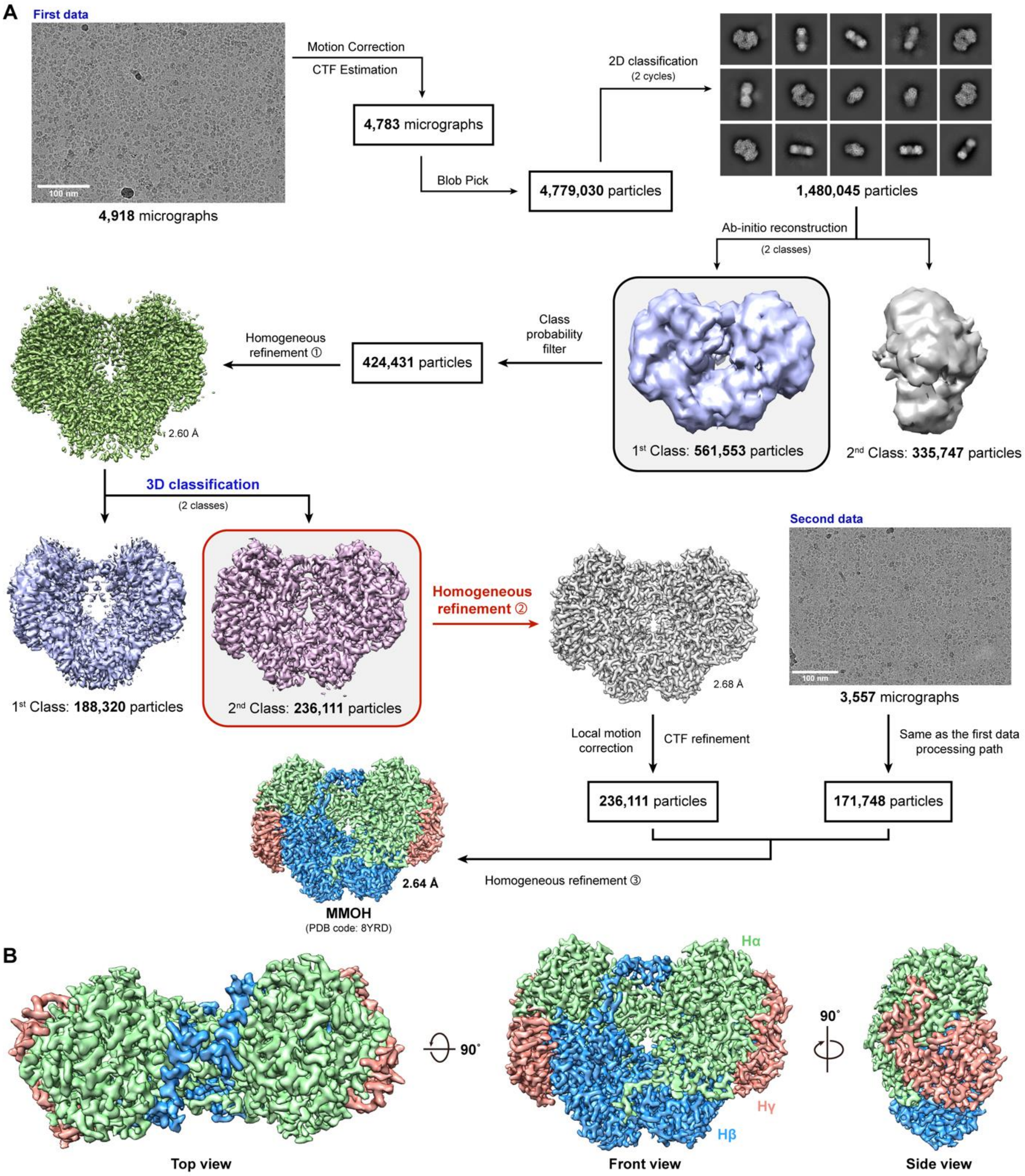


**Fig. S3. High flexibility like a "beating heart" of MMOH in native state with cryo-EM. a, Flow chart for cryo-EM image processing for MMOH (PDB: 8YRD).** The numbers next to the 3D map volumes indicate the resolution (unit: Å). As a result of data processing, the 3D map volume of MMOH was obtained with a 2.64 Å resolution. b, Top, front, and side views of MMOH (α-subunit, lime; β-subunit, marine; γ-subunit, deep salmon).

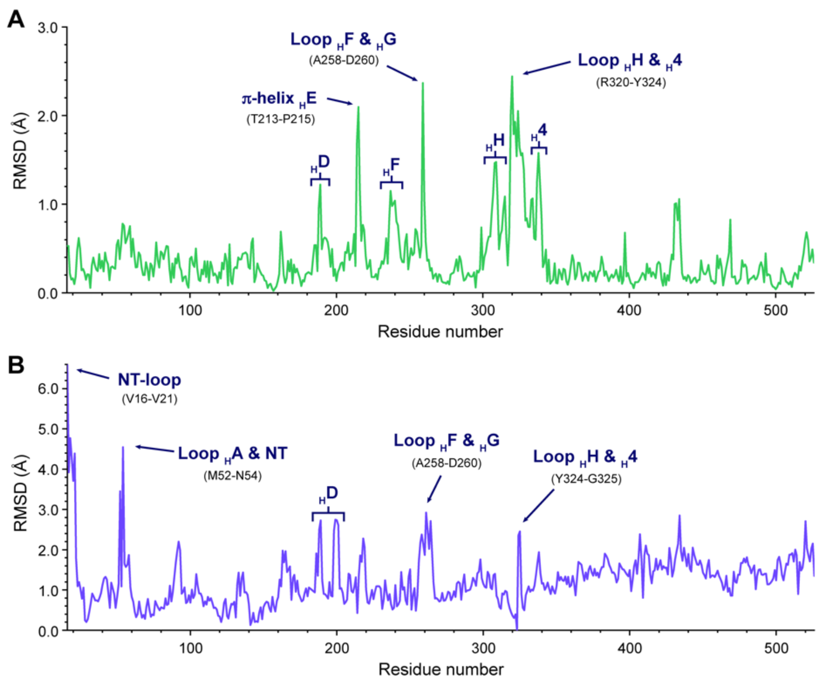


**Fig. S4. Comparison of conformational changes in MMOH α-subunit upon binding to MMOB.** (**A**-**B**) Root-mean-square deviation (RMSD) of the main chain ($C_\alpha$) per residue for the cryo-EM $HB^A$ with (**A**) cryo-EM $HB^B$ and (**B**) X-ray H-2B complex (PDB: 4GAM). Graph of RMSD per residue plotted using visual molecular dynamics (VMD, v.1.9.4a57) and the MultiSeq program.

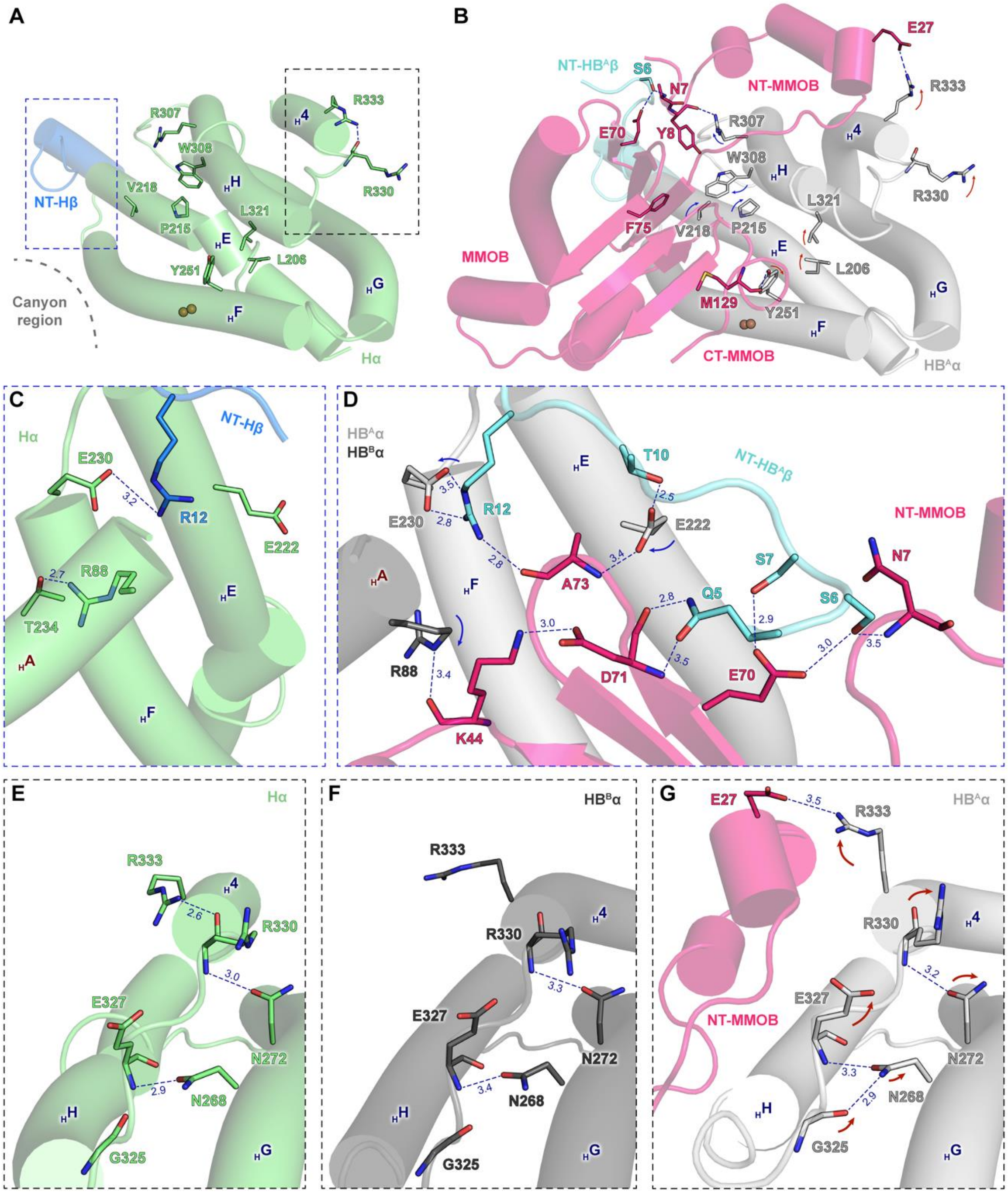


**Fig. S5. Helices that undergo significant conformational shifts upon binding of MMOB.** (**A**) cryo-EM MMOH (PDB: 8YRD). (**B**) cryo-EM HB$^A$ (PDB: 8XIW). The *N*-terminus (NT) of the MMOH β-subunit provides structural support, while the *C*-terminus (CT) and *N*-terminus (NT)-MMOB induce the elevation of the MMOH α-subunit. (**C**-**D**) Detailed molecular interactions at the canyon region mediated by the *N*-terminus of the MMOH β-subunit from (**C**) cryo-EM MMOH and (**D**) cryo-EM MMOH-1MMOB. A NT-HB$^A$β induces conformational changes in NT-MMOB, helices E and F in protomer A, and helix A in protomer B. (**E**-**G**) Detailed molecular interactions at the NT-MMOB with the MMOH α-subunit from (**E**) cryo-EM MMOH, (**F**) cryo-EM HB$^B$, and

(**G**) cryo-EM $HB^A$. The red arrows indicate conformational changes caused by NT-MMOB. The color of the arrows indicates conformational changes caused by NT- and CT-MMOB (red) and NT-$HB^A\beta$ (blue).

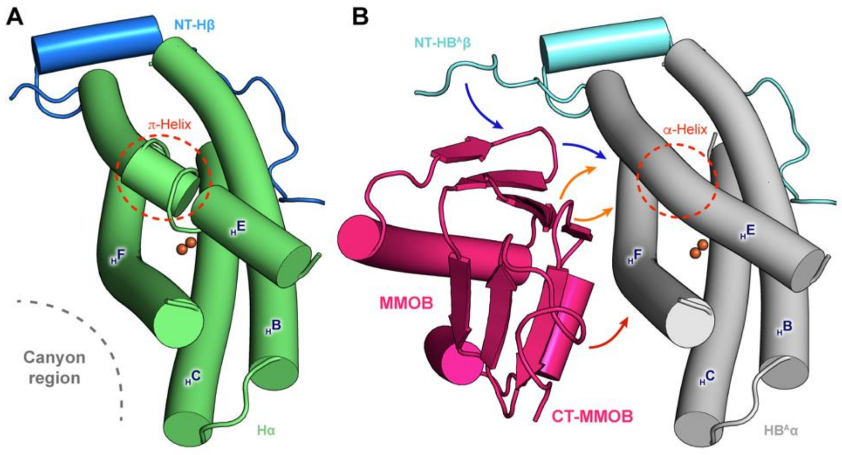


**Fig. S6. MMOB induced conformational changes of the four-helix bundles in the MMOH α-subunit from cryo-EM.** (**A**) MMOH. (**B**), cryo-electron microscopy (EM) $HB^A$. NT-MMOHβ provides structural support for helices B and C (*1, 2*). Due to the interaction of MMOB and NT-$HB^A$β with the MMOH canyon region, the π-helix of helix E is converted into an α-helix. The color of the arrows indicates conformational changes caused by CT-MMOB (red), NT-$HB^A$β (blue), and the core region of MMOB (orange).

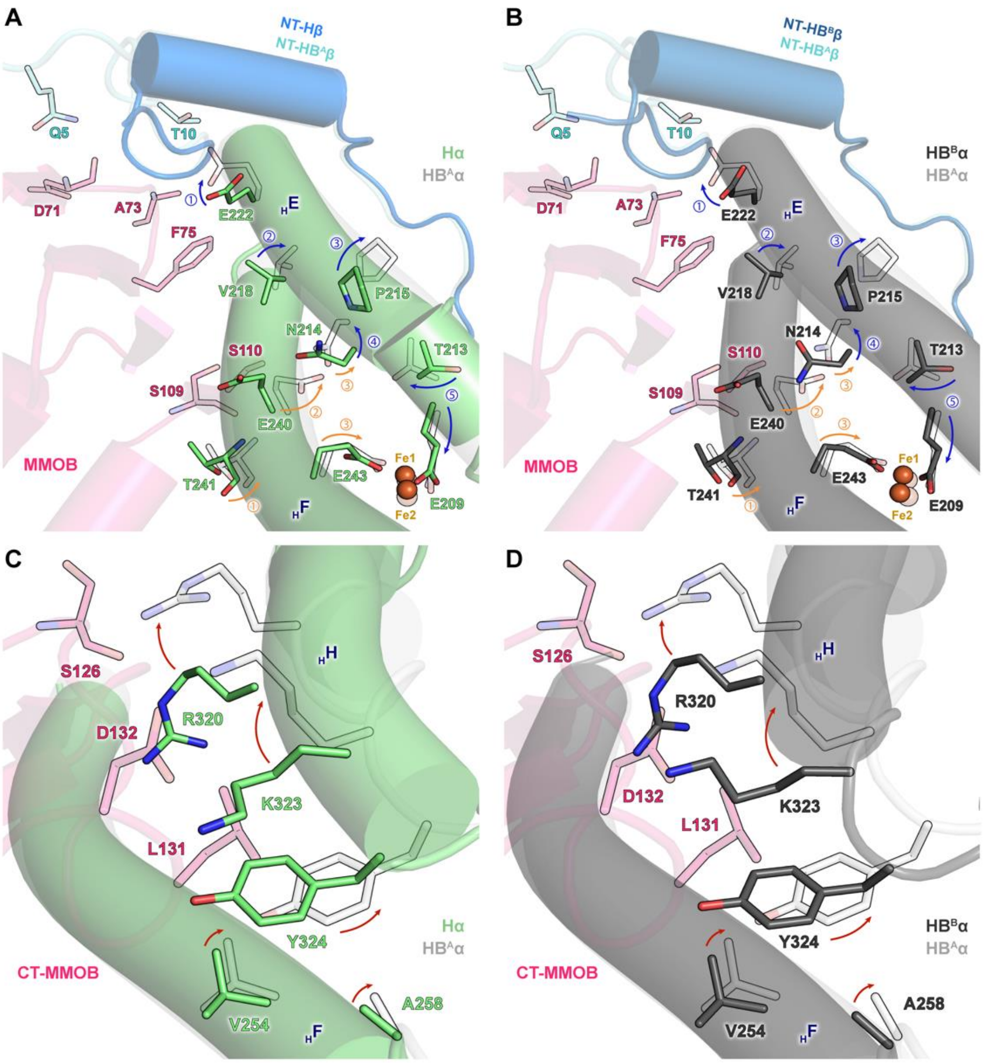


**Fig. S7. Conformational changes in MMOH induced by MMOB binding.** (**A** and **B**) Structural alignment of cryo-electron microscopy (EM) HB$^A$ with (**A**) cryo-EM MMOH and (**B**) cryo-EM HB$^B$ in the canyon region (*1*). c, d, Structural alignment of cryo-EM HB$^A$ with (**C**) cryo-EM MMOH and (**D**) cryo-EM HB$^B$ in C-terminal (CT)-MMOB with helices H and 4. CT-MMOB induces elevation of the MMOH α-subunit. The color of the arrows indicates conformational changes caused by CT-MMOB (red), NT-HB$^A$β (blue), and the core region of MMOB (orange).

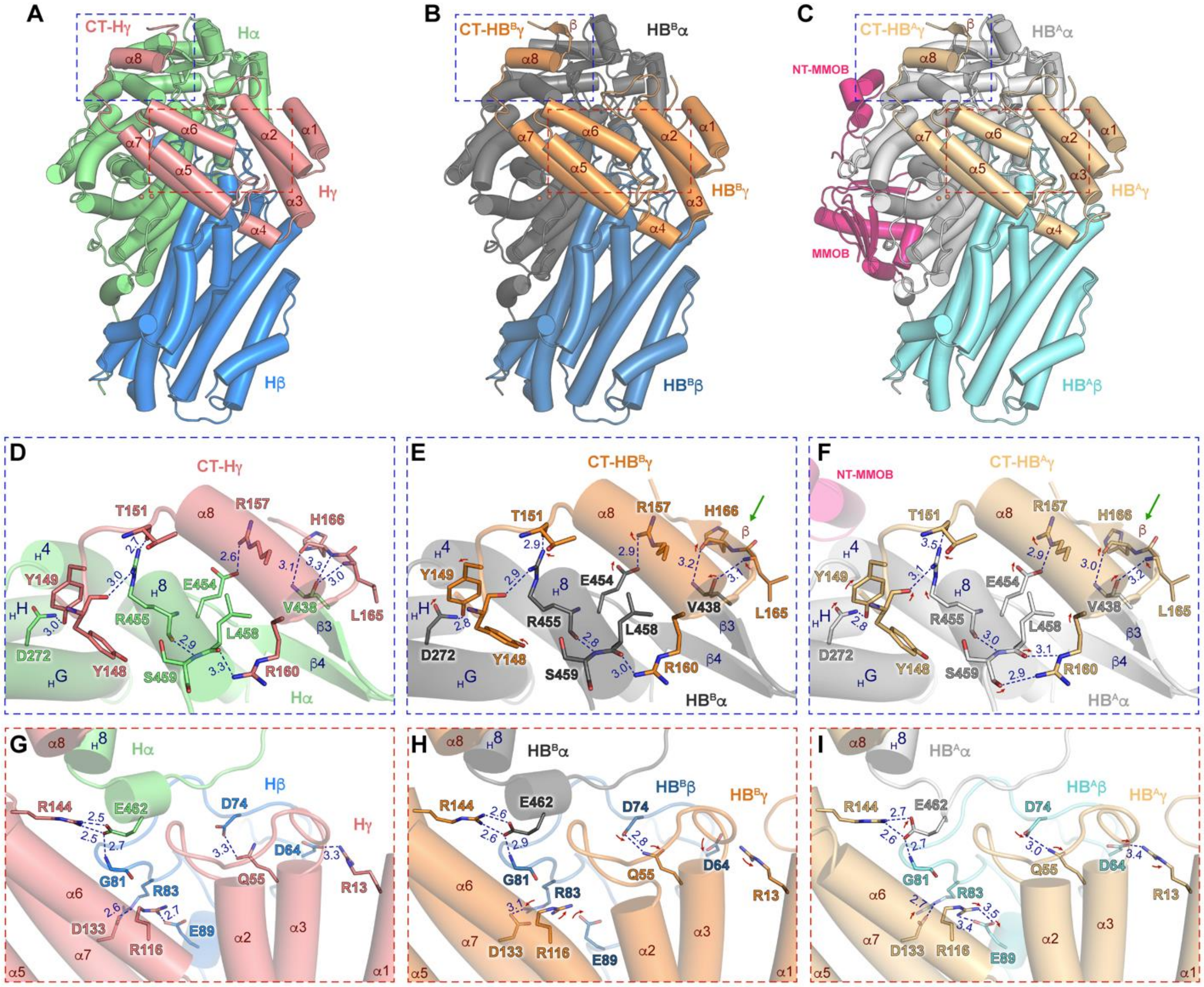


**Fig. S8. MMOH γ-subunit-mediated stabilization of conformational changes induced by MMOB binding.** (**A**-**C**) Overall structures of (**A**) cryo-EM MMOH (PDB: 8YRD), (**B**) cryo-EM HB$^{B}$ (PDB: 8XIW), and (**C**) HB$^{A}$ (PDB: 8XIW). The MMOH γ-subunit (MMOHγ) interacts with the MMOH α-subunit (MMOHα) and MMOH β-subunit (MMOHβ) to stabilize the overall structure. (**D**-**F**) The *C*-terminal region of MMOHγ interacts with MMOHα in (**D**) MMOH, (**E**) HB$^{B}$, and (**F**) HB$^{A}$. (**G**-**I**), The core region of MMOHγ interacts with MMOHβ in (**G**) cryo-EM MMOH, (**H**) HB$^{B}$, and (**I**) HB$^{A}$. Red arrows exhibit conformational changes caused by MMOB and green arrows indicate β-strand secondary structure of MMOHγ formed by MMOB binding.

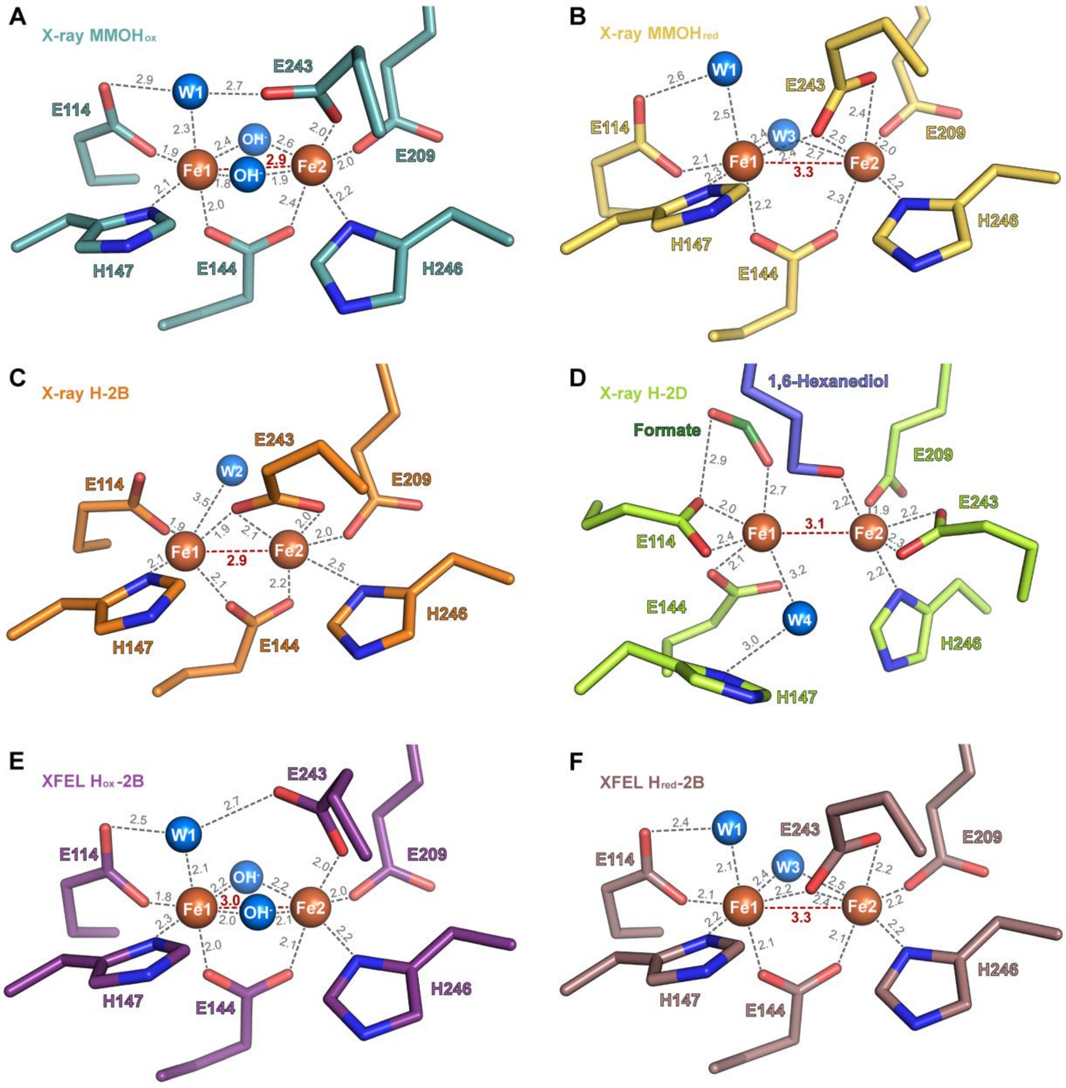


**Fig. S9. Di-iron active site in sMMO structure from X-ray crystallography and X-ray free electron laser (XEFL).** (**A**) X-ray, oxidized MMOH (PDB: 1MTY, resolution: 1.7 Å, teal). (**B**) X-ray, reduced MMOH (PDB: 1FYZ, resolution: 2.15 Å, yellow-orange). (**C**) X-ray, MMOH-2MMOB complex (PDB: 4GAM, resolution: 2.902 Å, tv_orange). (**D**) X-ray, MMOH-2MMOD complex (PDB: 6D7K, resolution: 2.6 Å, lemon). (**E**) XFEL, oxidized MMOH-2MMOB complex (PDB: 6YD0, resolution: 1.95 Å, violet-purple). (**F**) XFEL, reduced MMOH-2MMOB complex (PDB: 6YDI, resolution: 1.95 Å, dark-salmon). The two Fe atoms are coordinated by six residues, including four glutamates and two histidines. Water molecules are displayed as marine spheres and are numbered W1 and W4 according to their positions. The coordination distances are depicted in gray, and the Fe–Fe distances are represented in red. The unit of distance is Ångström (Å).

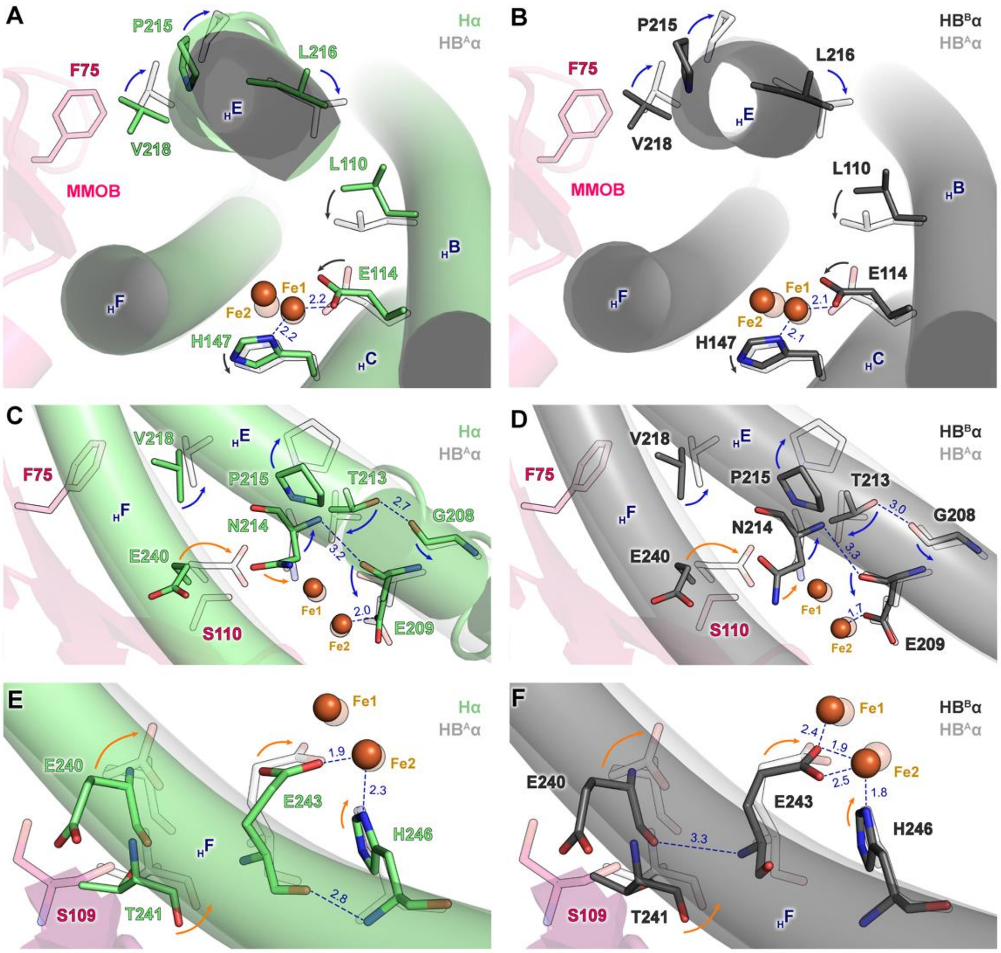


**Fig. S10. The key residues of the MMOH α-subunit related to the di-iron active site upon MMOB binding with cryo-electron microscopy (EM).** (**A** and **B**), Structural alignment of HB$^A$ with (**A**) MMOH and (**B**) HB$^B$ in the four-helix bundle. MMOB Phe75 triggers a shift in the residues coordinating Fe1 within the four-helix bundle (*3*). (**C** and **D**) Structural alignment of HB$^A$ with (**C**) MMOH and (**D**) HB$^B$ in helices E and F related to the π-helix and Fe1 (*4, 5*). (**E** and **F**), Structural alignment of HB$^A$ with (**E**) MMOH and (**F**) HB$^B$ in helix F related to residues coordinating with Fe2. The color of the arrows indicates conformational changes caused by NT-HB$^A$β (blue) and the core region of MMOB (orange). Black arrows indicate conformational changes in the residues of helices B and C. The color of the dotted box is based on the color of the arrow indicating the conformational changes.

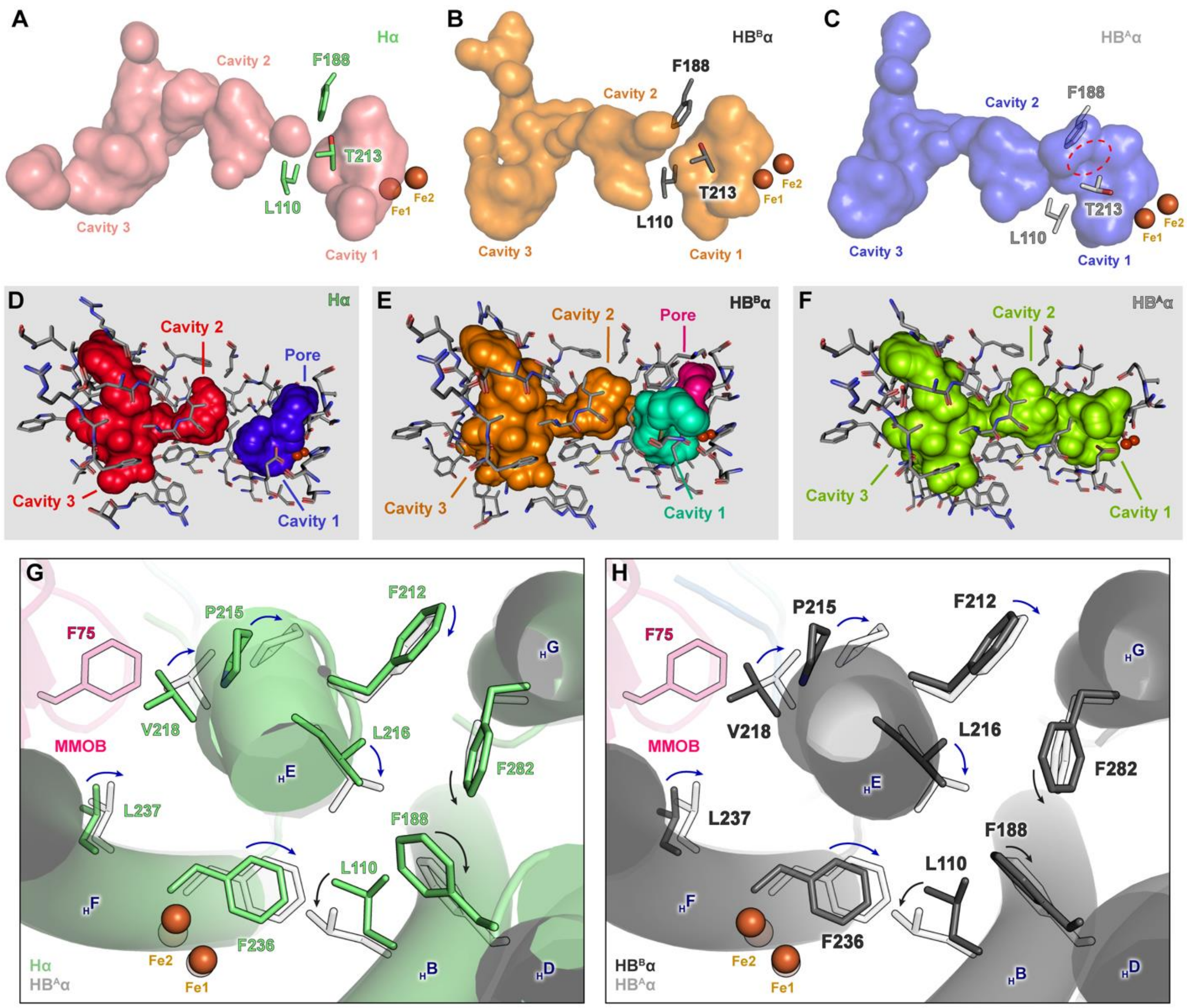


**Fig. S11. Analysis of cavities in cryo-EM of sMMO structures.** The cavities were displayed using (**A**-**C**) PyMOL 2.5.2 and (**D**-**F**) CAVER Analyst 2.0 BETA. Views of cavities 1, 2, and 3 are shown as translucent van der Waals surfaces in the interior of (**A**) MMOH (residue, lime; surface, deep-salmon), (**B**) $HB^B$ (residue, dark; surface, orange), and (**C**) $HB^A$ (residue, gray; surface, tv_blue). The cavities were displayed using PyMOL 2.5.2 with the following parameters: display quality, maximum; surface, cavities and pockets (culled); cavity detection radius, three solvent radii; cavity detection cutoff, five solvent radii; and HETATMs were ignored. The cavities were calculated in the interior of (**D**) MMOH, (**E**) $HB^B$, and (**F**) $HB^A$ using CAVER Analyst 2.0 BETA with the following parameters: probe, 1.5 Å; large probe, 3.0 Å. (**G**-**H**) Conformational changes in residues related to cavities upon binding MMOB with cryo-EM. Structural alignment of $HB^A$ with (**G**) MMOH and (**H**) $HB^B$. Blue arrows indicate conformational changes caused by NT-$HB^A\beta$. Black arrows indicate conformational changes in the residues of helices B, D, and G.

**Table S1. Cryo-EM data collection, refinement, and validation statistics.**

| | **MMOH** (PDB 8YRD) (EMD-39540) | **MMOH-1MMOB** (PDB 8XIW) (EMD-38391) |
|---|---|---|
| **Data collection and processing** | | |
| Magnification | 105,000 | 105,000 |
| Voltage (kV) | 300 | 300 |
| Electron exposure ($e^{-}/Å^2$) | 65.4 | 58.2 |
| Defocus range (μm) | -0.8 to -2.3 | -0.9 to -2.3 |
| Pixel size (Å) | 0.848 | 0.849 |
| Symmetry imposed | C2 | C1 |
| Micrographs collected/used | 8,475/8,171 | 15,007/9,471 |
| Initial particle images (no.) | 407,859 | 200,043 |
| Final particle images (no.) | 4,779,030 | 7,714,850 |
| Map resolution (Å) | 2.64 | 2.85 |
| Unmasked resolution at 0.5/0.143 FSC (Å) | 2.6/2.9 | 3.2/2.8 |
| Masked resolution at 0.5/0.143 FSC (Å) | 2.8/2.5 | 3.1/2.7 |
| **Structural refinement** | | |
| Initial model used (PDB code) | 1MTY | 4GAM |
| Model composition | | |
| Chains | 9 | 10 |
| Non-hydrogen atoms | 17,147 | 18,397 |
| Protein residues | 2,108 | 2,266 |
| Ligands | 4 | 4 |
| Mean *B* factors ($Å^2$) | | |
| Protein | 37.62 | 105.25 |
| Ligand | 47.98 | 120.74 |
| Water | 49.42 | 59.38 |
| R.m.s. deviations | | |
| Bond lengths (Å) (# > 4σ) | 0.005 (0) | 0.006 (0) |
| Bond angles (°) (# > 4σ) | 0.738 (0) | 1.053 (0) |
| Validation | | |
| Molprobity score | 0.86 | 0.69 |
| Clash score | 0.72 | 0.11 |
| Poor rotamers (%) | 0.92 | 0.86 |
| Cβ outliers (%) | 0.00 | 0.00 |
| CaBLAM outliers (%) | 0.29 | 0.63 |
| CC (mask) | 0.86 | 0.87 |
| Ramachandran plot | | |
| Favoured (%) | 97.33 | 97.25 |
| Allowed (%) | 2.67 | 2.75 |
| Disallowed (%) | 0.00 | 0.00 |

MMOH: methane monooxygenase hydroxylase
MMOB: MMO regulatory protein
FSC: fourier Shell Correlation
CaBLAM: Cα-based low-resolution annotation method

### Table S2. Metal coordination in the active sites of different forms of MMOH.

| Bond | Distance, Å | | | | | | | | |
|---|---|---|---|---|---|---|---|---|---|
| | Cryo-EM | | | X-ray crystallography | | | | XFEL | |
| | H | $HB^B$ | $HB^A$ | $H_{ox}$ | $H_{red}$ | H-2B | H-2D | $H_{ox}$-2B | $H_{red}$-2B |
| Fe1-Fe2 | 3.1 | 3.1 | 2.7 | 2.9 | 3.3 | 2.9 | 3.1 | 3.0 | 3.3 |
| Fe1-E114(OE1) | 2.2 | 2.1 | 1.9 | 1.9 | 2.1 | 1.9 | 2.0 | 1.8 | 2.1 |
| Fe1-E114(OE2) | $^a$ | $^a$ | $^a$ | $^a$ | $^a$ | $^a$ | 2.4 | $^a$ | $^a$ |
| Fe1-E144(OE1) | 2.0 | 2.3 | 1.9 | 2.0 | 2.2 | 2.1 | 2.1 | 2.0 | 2.1 |
| Fe2-E144(OE2) | 2.2 | 2.4 | 2.1 | 2.4 | 2.3 | 2.2 | $^a$ | 2.1 | 2.1 |
| Fe1-H147 | 2.3 | 2.1 | 2.5 | 2.1 | 2.3 | 2.1 | $^a$ | 2.3 | 2.2 |
| Fe2-E209 | 2.0 | 1.7 | 1.7 | 2.0 | 2.0 | 2.0 | 1.9 | 2.0 | 2.2 |
| Fe1-E243(OE1) | $^a$ | 2.4 | 2.2 | $^a$ | 2.4 | 1.9 | $^a$ | $^a$ | 2.2 |
| Fe2-E243(OE1) | $^a$ | 1.9 | 2.5 | $^a$ | 2.5 | 2.1 | 2.3 | $^a$ | 2.4 |
| Fe2-E243(OE2) | 1.9 | 2.5 | 2.5 | 2.0 | 2.4 | 2.0 | 2.2 | 2.0 | 2.2 |
| Fe2-H246 | 2.3 | 1.8 | 2.0 | 2.2 | 2.2 | 2.5 | 2.2 | 2.2 | 2.2 |
| W1-E114(OE2) | 2.7 | 2.7 | $^a$ | 2.9 | 2.6 | $^a$ | $^a$ | 2.5 | 2.4 |
| W2-E114(OE2) | $^a$ | $^a$ | 2.6 | $^a$ | $^a$ | $^a$ | $^a$ | $^a$ | $^a$ |
| W1-E243(OE1) | 3.5 | 2.9 | 2.5 | 2.7 | 3.1 | $^a$ | $^a$ | 2.7 | 2.9 |
| W1-Fe1 | 2.1 | 2.3 | 2.2 | 2.3 | 2.5 | $^a$ | $^a$ | 2.1 | 2.1 |
| W2-Fe2 | 3.9 | 4.0 | 4.0 | $^a$ | $^a$ | 3.5 | $^a$ | $^a$ | $^a$ |
| W3-Fe1 | $^a$ | $^a$ | $^a$ | $^a$ | 2.4 | $^a$ | $^a$ | $^a$ | 2.4 |
| W3-Fe2 | $^a$ | $^a$ | $^a$ | $^a$ | 2.7 | $^a$ | $^a$ | $^a$ | 2.5 |
| W4-Fe1 | $^a$ | $^a$ | $^a$ | $^a$ | $^a$ | $^a$ | 3.2 | $^a$ | $^a$ |
| W4-H147 | $^a$ | $^a$ | $^a$ | $^a$ | $^a$ | $^a$ | 3.0 | $^a$ | $^a$ |
| Fe1-$OH^-$ | $^a$ | $^a$ | $^a$ | 1.8/2.4 | $^a$ | $^a$ | $^a$ | 2.0/2.2 | $^a$ |
| Fe2-$OH^-$ | $^a$ | $^a$ | $^a$ | 1.9/2.6 | $^a$ | $^a$ | $^a$ | 2.1/2.2 | $^a$ |

$^a$Not applicable
MMOH: methane monooxygenase hydroxylase
Cryo-EM: cryogenic electron microscopy
XFEL: X-ray free electron laser

**References and Notes**


1. S. J. Lee, M. S. McCormick, S. J. Lippard, U.-S. Cho, Control of substrate access to the active site in methane monooxygenase. *Nature* **494**, 380-384 (2013).
2. H. Kim *et al.*, MMOD-induced structural changes of hydroxylase in soluble methane monooxygenase. *Sci. Adv.* **5**, eaax0059 (2019).
3. V. Srinivas, R. Banerjee, H. Lebrette, J. C. Jones, O. Aurelius, I.-S. Kim, C. C. Pham, S. Gul, K. D. Sutherlin, A. Bhowmick, J. John, E. Bozkurt, T. Fransson, P. Aller, A. Butryn, I. Bogacz, P. Simon, S. Keable, A. Britz, K. Tono, K. S. Kim, S.-Y. Park, S. J. Lee, J. Park, R. Alonso-Mori, F. D. Fuller, A. Batyuk, A. S. Brewster, U. Bergmann, N. K. Sauter, A. M. Orville, V. K. Yachandra, J. Yano, J. D. Lipscomb, J. Kern, M. Högbom, High-resolution XFEL structure of the soluble methane monooxygenase hydroxylase complex with its regulatory component at ambient temperature in two oxidation states. *J. Am. Chem. Soc.* **142**, 14249-14266 (2020).
4. D. Rinaldo, D. M. Philipp, S. J. Lippard, R. A. Friesner, Intermediates in dioxygen activation by methane monooxygenase: A QM/MM study. *J. Am. Chem. Soc.* **129**, 3135-3147 (2007).
5. J. C. Jones, R. Banerjee, K. Shi, H. Aihara, J. D. Lipscomb, Structural studies of the Methylosinus trichosporium OB3b soluble methane monooxygenase hydroxylase and regulatory component complex reveal a transient substrate tunnel. *Biochemistry* **59**, 2946-2961 (2020).
